# ImmunoPheno: A Computational Framework for Data-Driven Design and Analysis of Immunophenotyping Experiments

**DOI:** 10.64898/2026.02.01.703134

**Authors:** Lincoln Wu, Minh A. Nguyen, Zhangliang Yang, Sriya Potluri, Shamilene Sivagnanam, Nell Kirchberger, Apoorva Joshi, Kyung J. Ahn, Joseph S. Tumulty, Emylette Cruz Cabrera, Neil Romberg, Kai Tan, Lisa M. Coussens, Pablo G. Camara

## Abstract

Immunophenotyping is fundamental to characterizing tissue cellular composition, pathogenic processes, and immune infiltration, yet its accuracy and reproducibility remain constrained by heuristic antibody panel design and manual gating. Here, we present ImmunoPheno, an open-source computational platform that repurposes large-scale single-cell proteo-transcriptomic data to guide immunophenotyping experimental design and analysis. ImmunoPheno integrates existing datasets to automate the design of optimal antibody panels, gating strategies, and cell identity annotation. We used ImmunoPheno to construct a harmonized reference (HICAR) comprising 390 monoclonal antibodies and 93 human immune cell populations. Leveraging this resource, we algorithmically designed minimal panels to isolate rare populations, such as MAIT cells and pDCs, which we validated experimentally. We further demonstrate accurate cell identity annotation across publicly available and newly generated cytometry datasets spanning diverse technologies, including spatial platforms like CODEX. ImmunoPheno complements expert curation and supports continual expansion, providing a scalable framework to enhance the accuracy, reproducibility, and resolution of immunophenotyping.

## Introduction

Immunophenotyping — the use of antibody-based assays to identify and classify cells according to the antigens or proteins they express — has become a cornerstone of modern immunology^1^, with broad applications in oncology^2^ and stem cell biology^3,4^. Owing to its high throughput, scalability, and ability to isolate populations by complex phenotypes, flow cytometry has emerged as the dominant immunophenotyping technology, underpinning a vast body of biomedical research. More recently, spatially resolved technologies such as highly multiplexed quantitative immunohistochemistry (mIHC)^5,6^, imaging mass cytometry^7^, multiplexed ion beam imaging (MIBI)^8^, and co-detection by indexing (CODEX)^9^ have extended immunophenotyping to intact tissues, enabling the study of tissue architecture and cellular organization *in situ*.

Despite its utility and widespread adoption, immunophenotyping typically relies on manually defined gating strategies and predefined marker panels whose precision, efficiency, and reproducibility are often poorly characterized. Computational methods for clustering and dimensionality reduction partially address this limitation by grouping cells according to their antigenic profiles^10–13^. However, the resulting clusters still require manual annotation based on the presence or absence of specific protein combinations^14^. As a result, immunophenotyping remains a complex, subjective, and labor-intensive process, contributing to substantial variability in estimated cell population abundances derived from identical flow cytometry datasets across laboratories^15^.

These limitations have motivated sustained efforts to standardize immunophenotyping assays for common applications. Community-wide initiatives, including the Human ImmunoPhenotyping Consortium^16^ and the Immunoguiding Program^17^, have played central roles in establishing consensus marker sets and gating strategies. In parallel, computational approaches that leverage annotated reference cytometry datasets to transfer cell identity labels onto query datasets with overlapping antibody panels^18,19^, or to derive optimized gating strategies for isolating specific cell populations^20–22^, have enabled partial automation of immunophenotyping workflows. However, these approaches generally depend on user-provided reference datasets derived from the same tissue and using a substantially overlapping antibody panel, limiting their generalizability and often necessitating the generation of experiment-specific reference data.

Single-cell proteo-transcriptomic technologies, including CITE-seq^23^, Abseq^24^, and REAP-seq^25^, provide a powerful strategy for generating high-quality reference datasets by simultaneously profiling gene expression and surface protein markers in individual cells. These data enable high-resolution cell type and state annotations derived from transcriptomes to be directly linked to corresponding protein expression patterns measured with antibodies. In practice, however, the utility of this approach has been constrained by the size, scope, and fragmentation of available proteo-transcriptomic datasets^18,19^.

Here, we build upon this paradigm by developing computational methods that integrate information across the landscape of publicly available proteo-transcriptomic datasets, rather than relying on a single matched reference, to guide the design and analysis of conventional immunophenotyping experiments. The resulting open-source platform, ImmunoPheno, enables systematic, data-driven design of antibody panels and gating strategies, as well as automated annotation of high-resolution cell identities in immunophenotyping data. By explicitly accounting for heterogeneity across experiments and monoclonal antibodies, ImmunoPheno leverages existing single-cell resources to expand the effective reference space available for immunophenotyping analyses.

We anticipate that ImmunoPheno will serve as a broadly useful community resource, enabling the systematic repurposing of the vast but fragmented body of existing single-cell proteo-transcriptomic data to improve the resolution, accuracy, and reproducibility of immunophenotyping across laboratories and technologies.

## Results

### Overview of the ImmunoPheno platform

To streamline the design and analysis of immunophenotyping experiments using existing single-cell proteo-transcriptomic datasets, we developed ImmunoPheno, a computational platform comprising a server and a Python package (**Fig. 1**). The server hosts a relational database containing structured and harmonized data from published proteo-transcriptomic studies and provides an application programming interface (API) for programmatic access (**Supplementary Fig. 1**). The database includes normalized single-cell protein expression data together with curated cell type and state annotations derived from corresponding single-cell gene expression profiles. Antibodies, protein targets, cell identities, and tissues are indexed using unique identifiers from the Antibody Registry^26^, UniProtKB^27^, Cell Ontology^28^, and BRENDA tissue ontology^29^, respectively (**Fig. 1** and **Supplementary Fig. 1A**).

**Figure 1.**
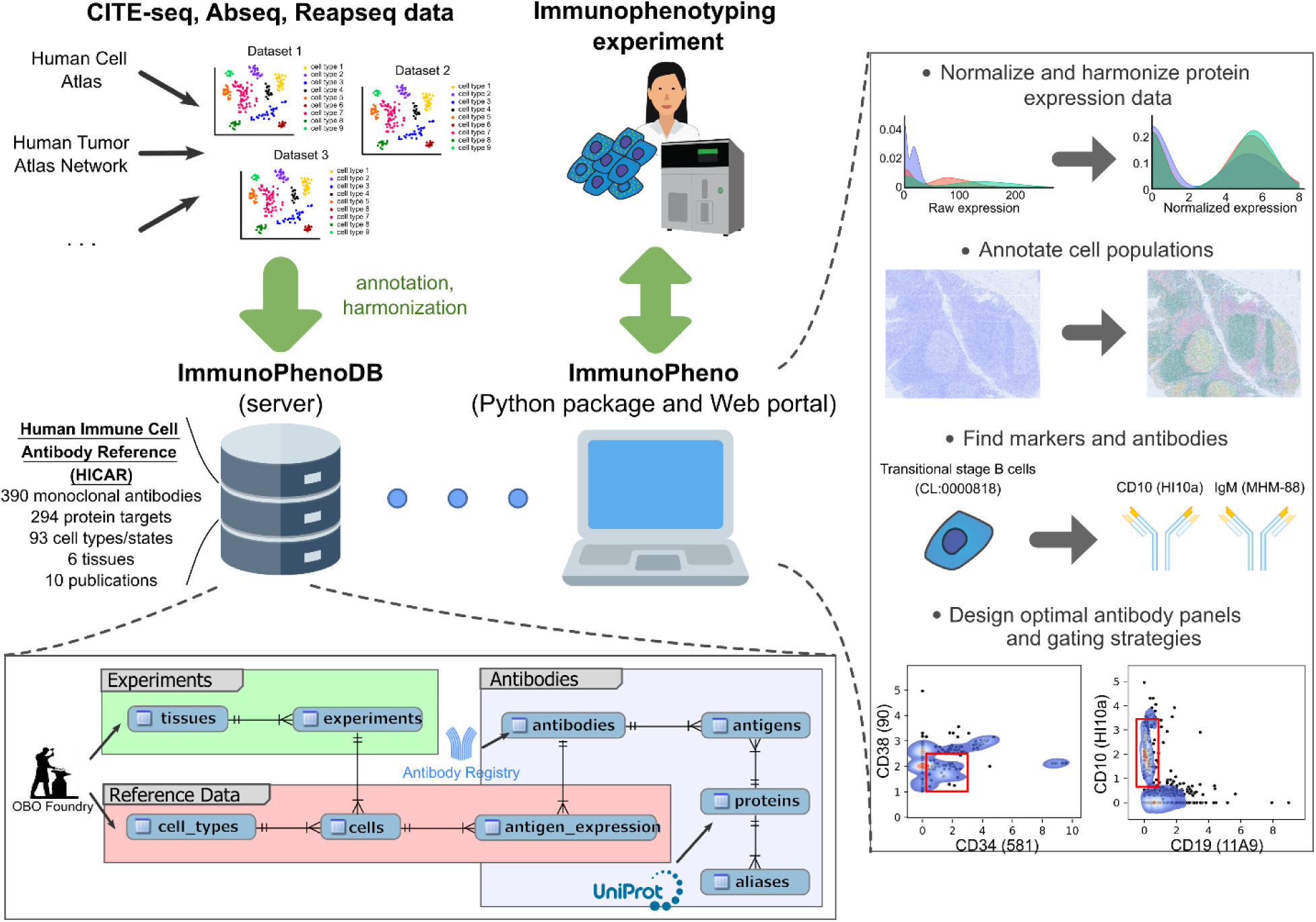
Overview of the ImmunoPheno platform. Schematic illustrating the different components of ImmunoPheno. Publicly available proteo-transcriptomic datasets are normalized, harmonized, annotated using OBO Foundry, UniProt, and Antibody Registry identifiers, and stored in the ImmunoPhenoDB relational database. Researchers generating or analyzing immunophenotyping data can then use the ImmunoPheno Python package or web portal to normalize and harmonize protein expression levels, automatically annotate individual cell identities, find antibodies with high specificity for a given population, and design optimal antibody panels based on data from previous experiments using those antibodies.

The Python package implements a suite of algorithms to enable data-driven analyses of immunophenotyping datasets using the reference data hosted on the server. These include methods for data normalization, automated cell identity annotation, and design of antibody panels tailored to specific phenotyping tasks. To account for variability between monoclonal antibodies targeting the same protein, ImmunoPheno operates at the level of individual monoclonal antibodies rather than protein targets. A key feature that distinguishes ImmunoPheno from existing data-driven approaches for cytometry data annotation and gating is its ability to integrate and leverage information across multiple reference datasets, thereby substantially expanding the effective reference space available for analysis.

ImmunoPheno is open-source and extensively documented. In addition to maintaining a central server (www.immunopheno.org) where users can contribute new reference datasets, we provide all the code and documentation for deploying independent servers. To facilitate accessibility, we also offer a web interface and an implementation of core functionalities within Galaxy^30^ (Methods). Together, these components make ImmunoPheno a valuable community resource for improving the phenotypic resolution, accuracy, and reproducibility of immunophenotyping experiments.

### Mixture-model-based harmonization improves cell state separation in immunophenotyping data

Widely used immunophenotyping technologies span a broad range of approaches, from conventional flow cytometry to highly multiplexed methods such as spectral flow cytometry^31^ and cytometry by time of flight (CyTOF)^32^, as well as spatially resolved platforms like quantitative mIHC^5,6^, imaging mass cytometry^7^, MIBI^8^, and CODEX^9^. These technologies differ substantially in their measurement modalities (sequencing read counts vs fluorescence intensity levels) and noise characteristics.

Mapping protein expression measurements from these diverse technologies onto reference single-cell proteo-transcriptomic datasets requires a normalization strategy that removes technology-specific background noise while preserving biologically meaningful signal. Building on our previous work^18^, we extended mixture-model-based normalization framework to improve background removal and accommodate a broader range of immunophenotyping technologies (Methods). Unlike existing batch-correction approaches for cytometry data^33–36^, this strategy enables new datasets to be incorporated into the reference database without requiring renormalization of preexisting data, facilitating scalability and efficient querying.

To evaluate the performance of this harmonization approach, we analyzed published CITE-seq^37^, CyTOF^12^, and CODEX^38^ datasets from human peripheral blood mononuclear cells (PBMCs), bone marrow mononuclear cells (BMMCs), and bone marrow tissue sections, respectively. The CITE-seq dataset included three replicates generated using different antibody concentrations^37^, whereas the CODEX dataset consisted of tissue sections from three different individuals, introducing substantial batch effects. Cell identities were annotated based on gene expression profiles using Azimuth^39^ (for CITE-seq data) or obtained from the original publications (for CyTOF and CODEX data), and each sample was normalized independently.

Before normalization, protein expression profiles exhibited substantial batch effects both within individual datasets and across technologies (**Fig. 2A**). These effects were markedly reduced after normalization, with protein expression distributions showing increased consistency across samples and platforms. Consistent with this observation, clustering of cells before normalization was largely driven by sample-of-origin rather than cell identity, resulting in extensive mixing of distinct cell types. After normalization, clustering was more strongly aligned with cell identity, reflecting improved separation of biologically meaningful cell states (**Fig. 2B**).

**Figure 2.**
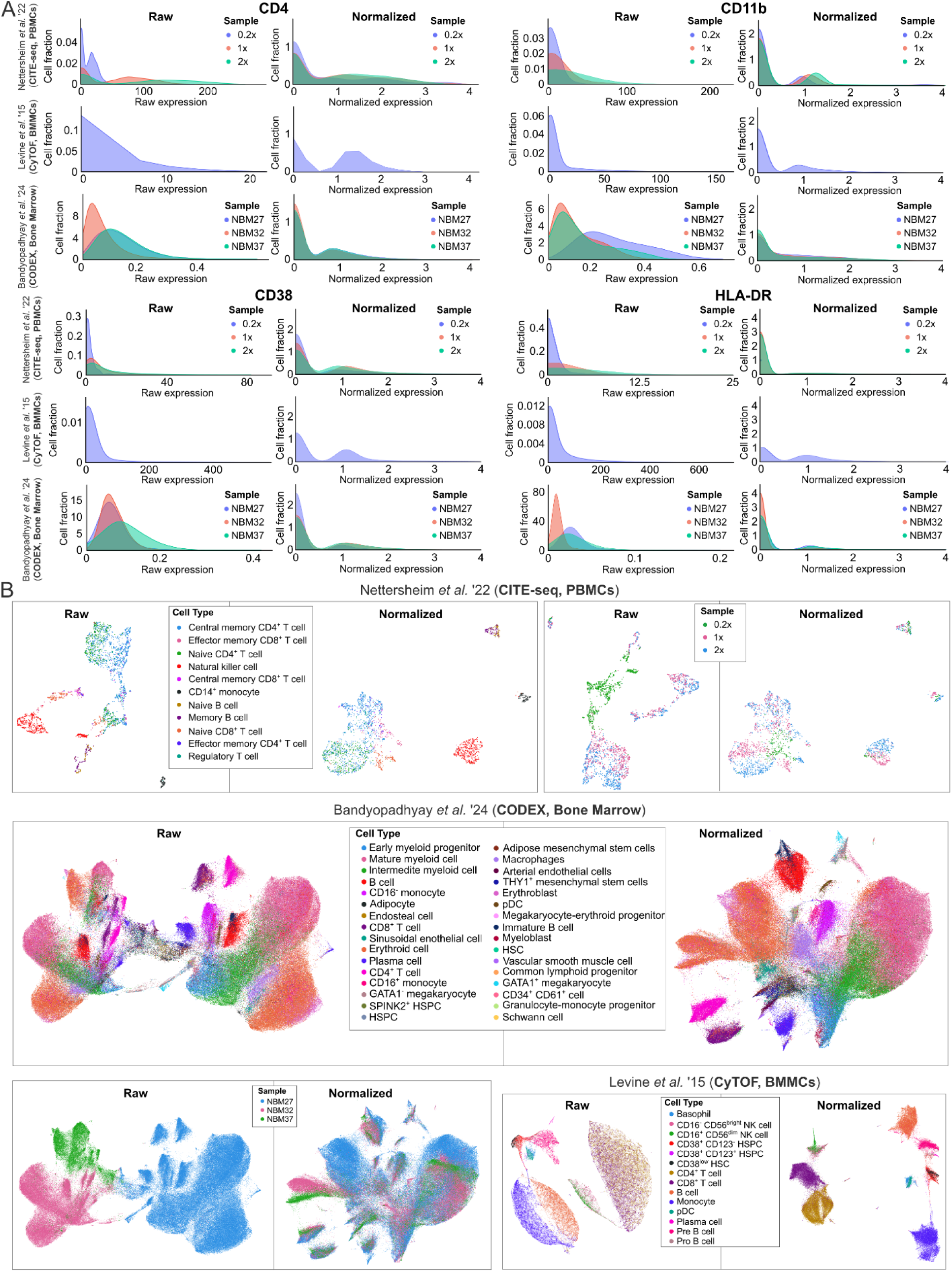
Harmonization of cytometry data across experiments and technologies. **A)** Distribution of raw and normalized protein expression values for four proteins across three datasets encompassing various cytometry technologies. Before normalization, the CITE-seq and CODEX datasets include substantial batch effects between samples. Each sample was normalized independently using Gaussian (for CyTOF and CODEX) and negative binomial (for CITE-seq) mixture models. The distribution of normalized expression values is consistent across samples and technologies. **B)** UMAP representations of the protein expression data before and after normalization. The representations are colored by the cell types identified from the transcriptomic data (for CITE-seq) or manual gating of the protein expression data (for CyTOF and CODEX). Additionally, the representations of the CITE-seq and CODEX datasets are also colored by the sample of origin. Before normalization, cells in the CODEX and CITE-seq datasets from the same cell type but from different samples appear in separate clusters. Additionally, several cell populations in the CyTOF data appear intermixed in the same cluster. These artifacts are substantially reduced or eliminated after normalization. PBMCs: peripheral blood mononuclear cells; BMMCs: bone marrow mononuclear cells; HSPC: hematopoietic stem and progenitor cells; HSC: hematopoietic stem cell; pDC: plasmacytoid dendritic cell.

These results demonstrate that mixture-model-based normalization applied independently to each sample effectively reduces batch effects and harmonizes protein expression measurements across diverse immunophenotyping technologies.

### A large-scale, harmonized proteo-transcriptomic resource of human immune cells

The advent of monoclonal antibodies and flow cytometry has transformed immunology into a field with a well-established taxonomic framework of cell types and states, defined by characteristic protein markers and standardized gating strategies. Accordingly, we focused on human immune cells to benchmark ImmunoPheno, enabling direct comparisons between cell annotations, gating strategies, and antibody panels generated by the platform and those established over decades of immunological research.

We applied mixture-model-based harmonization to 9 published proteo-transcriptomic datasets spanning 5 human immune tissues (**Supplementary Table 1**). To further expand tissue diversity, we generated a CITE-seq dataset from three human tonsils using a panel of 36 antibodies, and we processed it using the same pipeline (**Supplementary Fig. 2**). This tonsil dataset comprised 18,021 cells passing quality controls, with an average of 7,166 mRNA transcripts per cell. Collectively, the resulting resource, termed the Human Immune Cell Antibody Reference (HICAR), includes harmonized protein expression profiles for 115,979 cells across 6 human tissues, spanning 93 immune cell types annotated based on gene expression profiles, and covering 390 distinct monoclonal antibodies targeting 294 proteins (Methods) (**Fig. 3A**). We have uploaded HICAR to the ImmunoPheno server as reference data. To facilitate broader community use, we have developed a user-friendly web interface implementing a linear mixed-effects modeling framework to enable interactive data exploration and identification of antibodies associated with specific cell populations.

**Figure 3.**
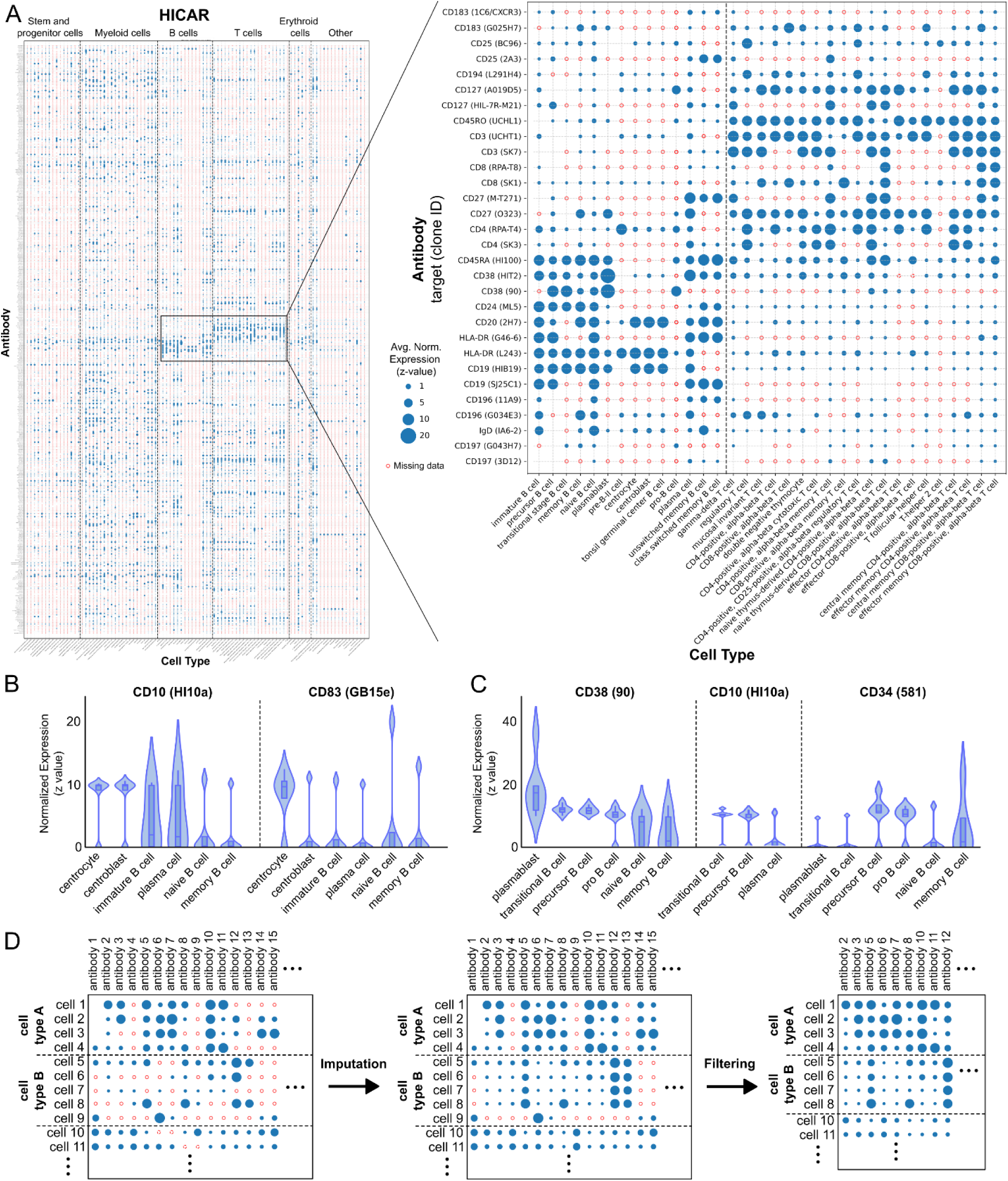
Human Immune Cell Antibody Reference (HICAR). **A)** Normalized protein expression data in the HICAR dataset, comprising 390 monoclonal antibodies and 93 cell populations. A section of the dataset corresponding to B cell and T cell populations and several monoclonal antibodies commonly used in the literature to gate these populations is zoomed in, showing consistency with known expression of cell population markers (for example, from the Human ImmunoPhenotyping Consortium^16^) and between monoclonal antibodies targeting the same protein. **B)** Normalized protein expression levels that best distinguish centrocytes and centroblasts from other B cell populations (anti-CD10) and from each other (anti-CD83) according to a linear mixed effects model applied to the HICAR data. **C)** Normalized protein expression levels that best distinguish transitional B cells from other B cell populations according to a linear mixed effects model applied to the HICAR data. **D)** Schematics of the imputation and filtering approach. Nearest-neighbor imputation is first applied within each cell type. A greedy approach is then used to remove the minimal amount of information (antibodies and cell types) required to eliminate all non-imputable data from the table.

We have validated HICAR through a wide range of queries, consistently recovering well-established markers and commonly used gating strategies, such as those recommended by the Human ImmunoPhenotyping Consortium^16^ (**Fig. 3A**). For example, querying antibodies that distinguish centroblasts and centrocytes from other B cell populations identified CD10 (clone HI10a) as a highly effective marker separating these populations from other B cells (Benjamini-Hochberg (BH) adjusted p-value < 10^-15^), and CD83 (clone GE15e) as a strong marker distinguishing centrocytes from centroblasts (BH adjusted p-value < 10^-15^) (**Fig. 3B**), consistent with previous reports^40,41^.

To test the utility of HICAR in challenging scenarios, we considered transitional B cells, an intermediate population between the pro B and mature stages of B cell development^42,43^. Due to their low abundance and transient nature, this population is relatively difficult to isolate, and several combinations of markers producing different outputs have been proposed, including CD19^+^ CD20^+^ CD10^+^ CD27^- 42,44,45^ and CD19^+^ CD38^+^ CD24^+^ CD27^- 42–44^. Querying HICAR for antibodies enriched in transitional B cells within the bone marrow B cell compartment identified CD38 (clone 90) and CD10 (clone H10a) as significantly upregulated markers (BH adjusted p-value < 10^-5^), detected in over 95% of transitional B cells. A subsequent query to identify other B cells in the bone marrow expressing these markers also revealed significant expression in pre- and pro-B cells (BH-adjusted p-value < 0.05) (**Fig. 3C**). We therefore performed a third query to identify markers that distinguish transitional B cells from pre- and pro-B cells, identifying CD34 (clone 581) as significantly downregulated in transitional B cells (BH-adjusted p-value < 10^-15^), being detected in >85% of pre- and pro-B cells but only in 7% of transitional B cells (**Fig. 3C**).

Thus, through three simple queries to HICAR, we found CD38^+^ CD10^+^ CD34^-^ as an effective gating strategy for isolating transitional B cells from other B cells in the bone marrow, illustrating the utility of HICAR for resolving challenging immunophenotyping tasks.

### Cross-dataset protein expression imputation and filtering enable integration across datasets

An important limitation of directly querying the ImmunoPheno database is the high degree of missing reference data. Each experiment in the database includes a different set of antibodies, and not all cell types or states are represented across every experiment. As a result, when HICAR is represented as a cell-by-antibody matrix, approximately 83% of the entries are missing (**Fig. 3A**).

To improve the robustness of ImmunoPheno’s inferences and enable integration across non-overlapping experiments, we implemented a k-nearest neighbor protein expression imputation strategy within each cell type (**Fig. 3D**) (Methods). Although several methods exist for protein expression imputation^46^, current approaches rely on computationally intensive integration of transcriptomic data from the same cells^39,47–49^ or learned transcriptome-proteome mappings^36,50,51^. We therefore developed a rapid, query-driven imputation approach that does not require paired transcriptomic measurements, allowing it to scale to large ImmunoPheno databases comprising harmonized protein expression data from dozens of reference datasets. We evaluated imputation accuracy by randomly masking entries in complete protein expression matrices from multiple datasets and comparing the imputed values with the original measurements. The imputed values showed strong concordance with the true values (**Supplementary Fig. 3A**), with Matthews Correlation Coefficient (MCC) values ranging from 0.77 to 0.97, depending on the proportion of missing data.

Although this imputation procedure substantially reduces data sparsity, it generally does not eliminate missing values entirely, as many entries are missing not at random (MNAR). For example, applying this approach to the HICAR dataset reduced the proportion of missing entries from 83% to 50%.

To remove residual missing values that could not be reliably imputed, we introduced a post-imputation filtering step that applies a greedy algorithm to iteratively remove antibodies and cells until a complete matrix with no missing entries is obtained (**Fig. 3D**; Methods). This filtering step prioritizes retention of antibodies and cell populations with broad and consistent coverage, minimizing the loss of informative features. To mitigate the risk of removing antibodies or cell populations that are sparsely represented across reference datasets but are important for the query dataset, the filtering algorithm explicitly balances the removal of cells versus antibodies through a tunable parameter, allowing users to prioritize retention of rare populations or broader antibody coverage depending on the analysis goal.

To evaluate the efficiency of post-imputation filtering, we simulated MNAR conditions by masking all entries of one or more antibodies in one or more cell types and quantified the ratio of MNAR entries to the total number of entries removed by the algorithm. Across these simulations, post-imputation filtering remained relatively efficient even under substantial MNAR conditions, with efficiencies typically ranging from 45% to 70% when 40% of the data was MNAR (**Supplementary Fig. 3B, C**).

Together, this imputation and filtering framework enables robust cross-dataset integration of information within ImmunoPheno, facilitating downstream applications such as antibody panel design and cell identity annotation in immunophenotyping analyses.

### Design of optimal antibody panels using ImmunoPheno

Having established an informatics infrastructure to harmonize, integrate, impute, and query published proteo-transcriptomic data, we next developed algorithms for the data-driven design and analysis of immunophenotyping experiments using the reference data stored in the ImmunoPheno database.

The design of antibody panels and gating strategies for isolating cell populations has traditionally relied on qualitative or partially subjective decisions informed by investigator experience. Although several efforts have sought to standardize cytometry assays in immunology^1,16,17,52^, systematically exploring the vast combinatorial space of possible antibody panels remains intractable without algorithmic approaches. For example, using the antibodies cataloged in the HICAR database, there are more than 2 × 10^19^ distinct 10-antibody panels and 6 × 10^44^ distinct 30-antibody panels that can be formed.

To enable data-driven design of immunophenotyping antibody panels, we coupled the imputation approach described above with a Classification and Regression Tree (CART) model^53^ to identify combinations of *k* antibodies that distinguish a user-defined set of target cell populations from background populations (Methods). In addition to identifying an optimal antibody set, the CART model generates corresponding gating strategies for isolating each target population, together with estimates of expected purity (precision) and yield (recall).

We first tested this approach by generating antibody panels for isolating commonly studied immune cell populations based on HICAR. The resulting panels were consistent with widely used and independently validated markers and gating strategies^16^. For example, a query for a four-antibody panel to distinguish T cell subsets in PBMCs produced gating strategies for isolating CD4^+^ and CD8^+^ naïve and memory T cells using anti CD4 (clone RPA-T4), CD8 (clone RPA-T8), CD2 (clone RPA-2.10), and CD27 (clone M-T271) antibodies, with expected purity and yield exceeding 90% for most subsets (**Figs. 4A, B**). Similarly, a query for a two-antibody panel designed to separate naïve and memory B cells within the B cell compartment identified CD44 (clone BJ18) and CD72 (clone 3F3), again achieving expected purity and yield above 90% (**Fig. 4C**). These results demonstrate that ImmunoPheno recapitulates established immunophenotyping strategies using HICAR data.

**Figure 4.**
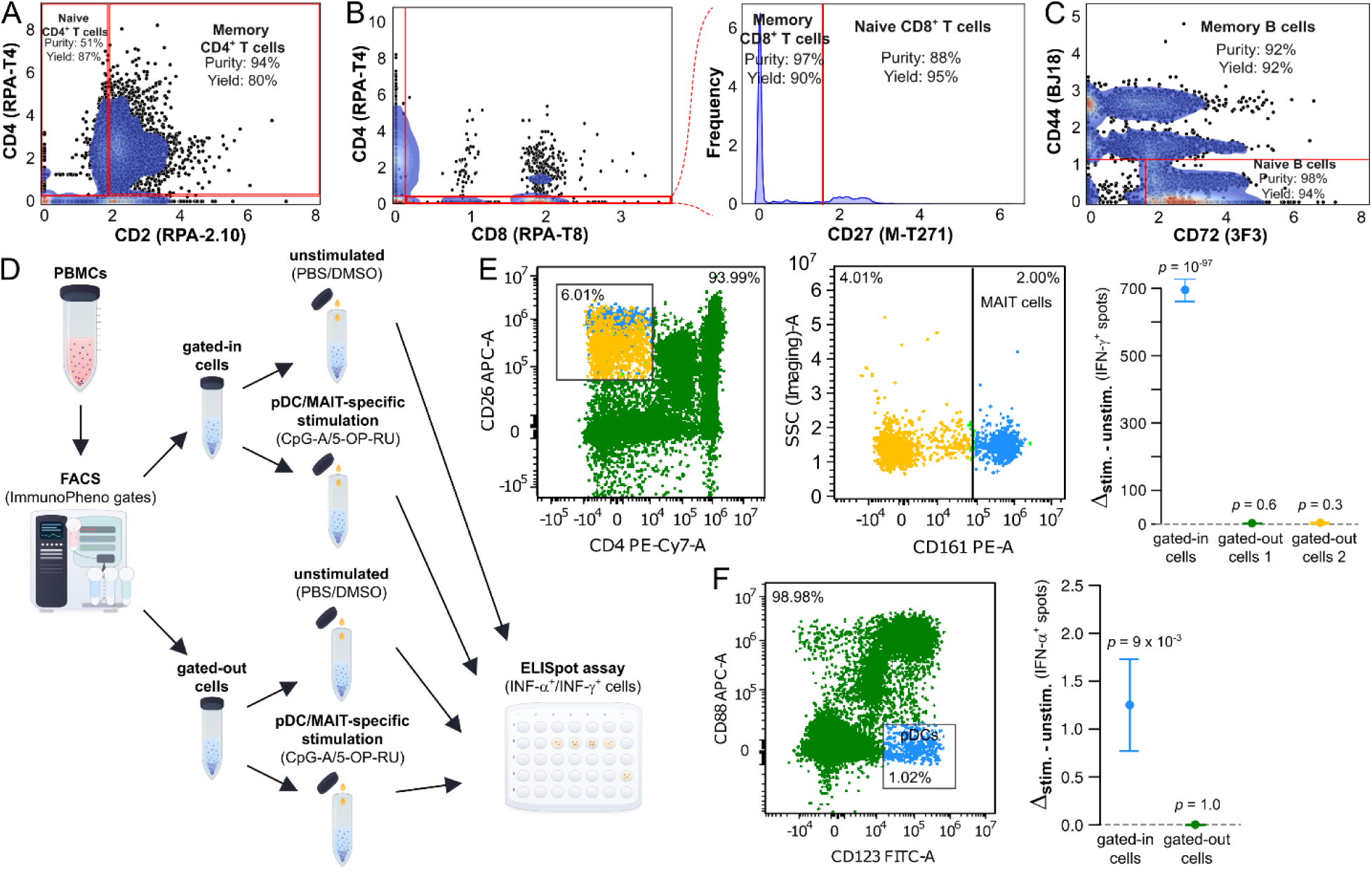
Automated design of antibody panels and gating strategies. **A, B)** Four-antibody panel and gating strategies generated by ImmunoPheno using HICAR data for profiling T-cell subsets in PBMCs. **C)** Two-antibody panel with gating strategy and expected yield and purity for isolating naïve and memory B cells from a B cell sample. **D)** Schematic of functional assay used to test ImmunoPheno-generated antibody panels and gating strategies for isolating MAIT cells and pDCs from PBMCs. **E)** FACS implementation of ImmunoPheno gates for isolating MAIT cells using a three-antibody panel, and results of functional assay showing a significant increase in IFN-γ^+^ cells upon 5-OP-RU stimulation in gated-in cells but not gated-out cells. Error bars indicate standard errors. **F)** FACS implementation of ImmunoPheno gates for isolating pDCs using a three-antibody panel, and results of functional assay showing a significant increase in IFN-α^+^ cells upon CpG-A stimulation in gated-in cells but not gated-out cells. Error bars indicate standard errors.

We next assessed whether ImmunoPheno could design minimal antibody panels to isolate rare immune cell populations from human PBMCs that could be readily incorporated into larger, more comprehensive panels. Specifically, we generated three- and two-antibody panels for isolating mucosal-associated invariant T (MAIT) cells and plasmacytoid dendritic cells (pDCs), respectively. MAIT cells constitute approximately 1–10% of PBMCs in healthy adults and are typically isolated using 5-OP-RU/MR1 tetramers^54^ (e.g., from the NIH Tetramer Core Facility) and CD3 co-staining. Similarly, pDCs represent less than 1% of PBMCs and are commonly isolated by magnetic bead-based negative selection followed by flow cytometric enrichment based on CD123 and CD303/CD304 expression^55^. Instead, the gating strategies produced by ImmunoPheno were based on the marker combinations CD4^-^ CD26^+^ CD161^+^ for MAIT cells and CD88^-^ CD123^+^ for pDCs (**Supplementary Fig. 4**).

To experimentally test these strategies, we implemented the associated gating schemes by fluorescence-activated cell sorting (FACS) and performed functional evaluation of the sorted populations using an enzyme-linked immunospot (ELISpot) assay (**Fig. 4D** and **Supplementary Table 2**). For MAIT cells, gated-in and gated-out populations were stimulated with 5-OP-RU, a synthetic ligand that specifically activates MAIT cells through major histocompatibility complex class I-related (MR1) presentation^56^, and interferon-γ (IFN-γ) production was measured by ELISpot. For pDCs, gated-in and gated-out populations were stimulated with the class A CpG oligonucleotide ODN 2216, a potent Toll-like receptor 9 (TLR9) agonist^57^, followed by measurement of interferon-α (IFN-α) production.

Consistent with successful isolation of MAIT cells and pDCs, we observed a marked enrichment of IFN-γ^+^ or IFN-α^+^ cells in the gated-in populations following specific stimulation, whereas the gated-out populations showed no detectable increase in IFN-γ/α-producing cells (**Fig. 4E, F**).

Collectively, these results demonstrate that ImmunoPheno, coupled with HICAR reference data, enables the data-driven design of effective antibody panels and gating strategies for isolating both common and rare immune cell populations.

### Automated cell identity annotation using ImmunoPheno

Computational approaches that transfer transcriptome-derived cell identity annotations from proteo-transcriptomic reference datasets onto cytometry data have substantially improved the resolution and reproducibility of immunophenotyping analyses^18,19^. However, existing methods require extensive overlap between the antibody panels used in the query and reference datasets, often necessitating the generation of experiment-specific reference data.

We reasoned that the imputation framework implemented in ImmunoPheno could overcome these limitations by integrating reference data from multiple datasets, thereby enabling greater flexibility in experimental design. To test this hypothesis, we augmented Spatial Transcriptomics via Epitope Anchoring (STvEA)^18^ with methods for filtering low-quality annotations based on label uncertainty and consistency of protein expression profiles among cells assigned the same label, and integrated the resulting framework into ImmunoPheno (Methods).

To evaluate the accuracy of automated cell annotations from ImmunoPheno, we analyzed two published CITE-seq datasets of PBMCs and bone marrow cells^58,59^, containing 42 and 22 antibodies, respectively. In each case, single-cell protein expression data were used as the query, while transcriptome-derived cell identities served as ground truth. Annotation performance was assessed both at the population level, based on recovery of transcriptomically defined cell populations, and at the single-cell level using the MCC.

As an upper bound on achievable performance, we first annotated each dataset using itself as the reference. Based solely on protein expression profiles, ImmunoPheno successfully recovered 17 of 20 and 20 of 24 transcriptomic cell populations in the PBMC and bone marrow datasets, respectively (**Supplementary Fig. 5**). At the single-cell level, the protein-based annotations showed strong concordance with the transcriptome-derived labels, with an MCC of 0.8 (**Fig. 5A**).

**Figure 5.**
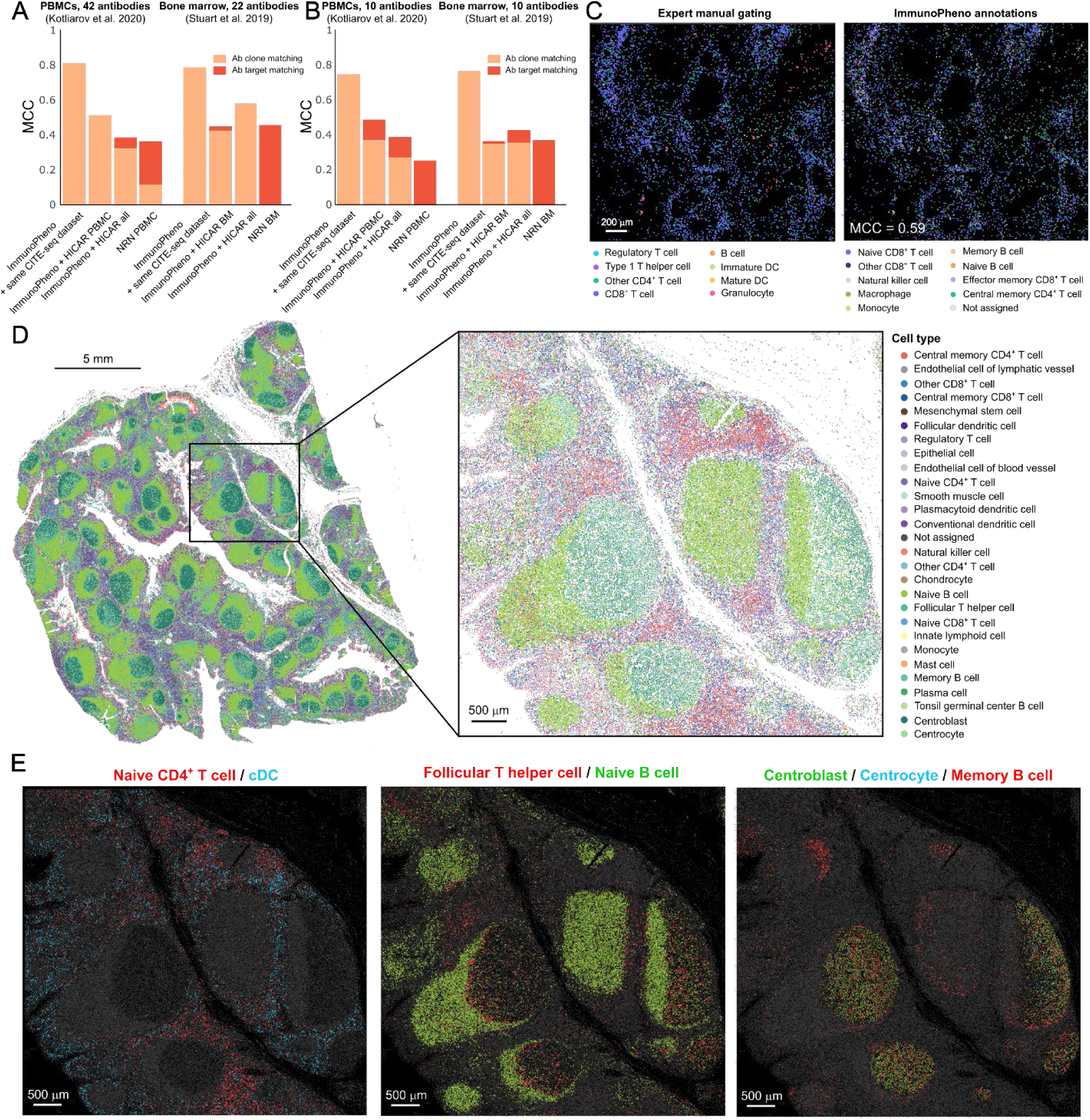
Automated cell identity annotation. **A, B)** Matthews correlation coefficient (MCC) of single-cell identity annotations produced by ImmunoPheno and NRN^19^ for two published PBMC and bone marrow CITE-seq datasets^58,59^. Results are shown using either the complete antibody panels (A) or a reduced panel of 10 antibodies (B). Performance is reported for cases in which antibodies in the query and reference were matched by clone ID or by protein target. **C)** Pancreatic ductal adenocarcinoma tumor profiled by chromogenic mIHC using a 22-antibody panel. Tumor-infiltrating CD45^+^ cells were annotated manually using established gating strategies (left) and with ImmunoPheno (right), based on 12 antibodies with matched targets in HICAR. **(D, E)** Automated annotation of cell identities in CODEX data from a human tonsil. **D)** Tissue section profiled with CODEX using a 50-antibody panel, with cell identities automatically annotated by ImmunoPheno based on HICAR, identifying 27 distinct cell types. **E)** Magnified region highlighting the spatial organization of cell populations involved in different stages of antigen presentation and B-cell activation.

We next annotated each dataset using HICAR as the reference, restricted to datasets derived from the same tissue type as the query and excluding the query dataset itself. Under these conditions, ImmunoPheno recovered 15 of 20 transcriptomic cell populations in the PBMC dataset, achieving a single-cell MCC of 0.52 (**Fig. 5A** and **Supplementary Fig. 5**). In contrast, when applied to the bone marrow dataset, only 7 of 24 populations were recovered, with a single-cell MCC of 0.37. We reasoned that this reduced performance likely reflected the limited representation of bone marrow data within the HICAR database. Consistent with this interpretation, repeating the analysis using the full HICAR database (while still excluding the query dataset) improved performance, enabling recovery of 13 of 24 transcriptomic cell populations and increasing the single-cell MCC to 0.58 (**Fig. 5A** and **Supplementary Fig. 5**).

To assess the impact of antibody panel size on annotation accuracy, we next repeated these analyses using only a subset of 10 antibodies from each query dataset. Across both datasets, MCC values were moderately reduced relative to those obtained with the full antibody panels (**Fig. 5B**), indicating that ImmunoPheno remains robust to relatively small antibody panels. Moreover, a substantial fraction of the performance loss could be mitigated by allowing antibody matching between query and reference data based on protein targets rather than antibody clones (**Fig. 5B**).

Finally, to place these results in the context of existing annotation methods, we evaluated whether ImmunoPheno’s ability to integrate reference data from multiple sources provides an advantage over approaches that rely on a single large-scale proteo-transcriptomic reference dataset. To this end, we annotated the PBMC and bone marrow query datasets using the Nearest Rank Neighbors (NRN) method and the large-scale PBMC and bone marrow single-cell proteo-transcriptomic reference datasets introduced by Triana *et al.*^19^, which comprise a panel of 197 antibodies. In this analysis, NRN recovered 13 and 12 transcriptomic populations in the PBMC and bone marrow datasets, respectively, with single-cell MCCs of 0.25 and 0.48 (**Figs. 5A, B**, and **Supplementary Fig. 6**), lower than those achieved by ImmunoPheno using the HICAR reference.

Together, these results demonstrate the value of leveraging multiple reference datasets for automated cell identity annotation in immunophenotyping data, particularly in settings where the antibody overlap with individual reference datasets is limited or where no single comprehensive, tissue-matched reference dataset is available. Moreover, they suggest practical strategies for applying ImmunoPheno: when the database contains multiple datasets from the same tissue type as the query, restricting annotation to those references yields optimal performance, whereas incorporating datasets from related tissues provides an effective alternative when tissue-matched data are sparse.

### ImmunoPheno cell annotations are consistent with manual expert annotations

To further evaluate the performance of ImmunoPheno’s automated annotations in complex tissue settings, we analyzed chromogenic mIHC profiles from two pancreatic ductal adenocarcinoma (PDAC) tumors using a 22-antibody panel comprising both intracellular and surface markers^60^. Of these 22 antibodies, only one was directly present in HICAR, whereas 11 others targeted proteins for which alternative antibodies were represented in HICAR. We selected two regions of interest with abundant immune infiltration and manually annotated 7 immune cell populations using established gating strategies^60^ (**Supplementary Fig. 7A, B**). These manual annotations were then compared with those generated by ImmunoPheno using the 12 antibodies with matched targets in HICAR (**Fig. 5C** and **Supplementary Fig. 7C**).

ImmunoPheno successfully annotated 77% and 80% of the cells identified by manual gating in the two sections, with single-cell MCCs of 0.59 and 0.54. Notably, in several cases the automated annotations achieved finer resolution, distinguishing 9 and 10 immune cell populations in the two sections, respectively. In addition, some CD45^+^ cells that were left unannotated in the manual analysis received annotations from ImmunoPheno.

The main discrepancies between ImmunoPheno and manual annotations arose from two sources (**Supplementary Fig. 7D**). Granulocytes and dendritic cells were frequently misclassified or unannotated due to their absence from the HICAR database, as these cell populations are typically depleted from CITE-seq data. In addition, regulatory T cells could not be resolved using surface markers alone and were annotated by experts using the intracellular markers FOXP3.

Despite these localized discrepancies arising from the limited overlap with HICAR, the overall concordance between ImmunoPheno and expert manual annotations was high, demonstrating the robustness and utility of ImmunoPheno in complex contexts such as solid tumors with very limited antibody panel overlap.

### Annotation of multiplexed immunohistochemistry data from a human tonsil

Finally, to demonstrate the utility of ImmunoPheno for large-scale analysis of human mIHC data, we profiled a human tonsil tissue section using CODEX with a 50-antibody panel, 36 of which were represented in HICAR (**Supplementary Table 3**). Applying ImmunoPheno to this dataset, we annotated 1,063,380 individual cells that passed all quality controls, corresponding to 27 distinct cell types and 73% of the total segmented cells (**Fig. 5D**).

ImmunoPheno cell annotations accurately recapitulated the known microanatomy of the tonsil, consisting of numerous lymphoid follicles containing germinal centers rich in proliferating cells surrounded by mantle zones; diffuse lymphatic tissue populated by T cells occupying the interfollicular regions; and a stratified squamous epithelium covering the surface.

The spatial distribution of annotated cell populations also reflected the characteristic dynamics of antigen presentation and B-cell activation in the tonsil, with dendritic cells capturing antigens in the subepithelial region and presenting them to naïve CD4^+^ T cells within the interfollicular zones (**Fig. 5E**). The subsequent interaction of activated T cells with naïve B cells in adjacent follicles was also captured, as well as the sequential differentiation of B cells from centroblasts to centrocytes before giving rise to memory B cells (**Fig. 5E**). These and other local cell-cell interactions were also recapitulated through quantitative cellular neighborhood analysis of the annotations (**Supplementary Fig. 8**).

Altogether, these results highlight the phenotypic resolution and biological interpretability of ImmunoPheno’s automated cell identity annotations and underscore its potential to reveal spatially organized immune processes in complex human tissues.

## Discussion

The emergence of massively parallel single-cell RNA-seq technologies has produced an explosion of transcriptomic data from mammalian tissues, including an increasing number of single-cell proteo-transcriptomic datasets with high-resolution annotations of cell types and states. Repurposing these data to guide the design and analysis of immunophenotyping experiments offers a powerful strategy to improve their phenotypic resolution, accuracy, and reproducibility.

To support this approach, we developed ImmunoPheno, a computational platform that provides end-to-end tools for constructing and querying structured databases of harmonized proteo-transcriptomic reference data from multiple sources, and designing antibody panels, identifying efficient gating strategies, and annotating individual cell identities based on these data. We demonstrated the platform by creating HICAR, a resource comprising proteo-transcriptomic data for 390 monoclonal antibodies and 93 human immune cell populations derived from 10 datasets. Using HICAR and ImmunoPheno, we designed small antibody panels and gating strategies to isolate both common and rare immune cell populations, and annotated cell identities in publicly available and newly generated cytometry datasets spanning multiple technologies, including spatially resolved platforms. Across these settings, ImmunoPheno matched manually curated antibody panels, gating strategies, and cell annotations that reflect decades of immunological expertise.

ImmunoPheno differs from existing approaches in several key respects. First, it is designed to integrate information across multiple heterogeneous proteo-transcriptomic datasets rather than relying on a single reference, substantially expanding the effective reference space available for analysis. Second, it operates at the level of individual monoclonal antibodies, explicitly accounting for clone-specific differences in protein detection that are typically ignored in protein-centric annotation frameworks. Third, beyond automated cell identity annotation, ImmunoPheno directly addresses the upstream experimental design problem by enabling data-driven selection of antibody panels and corresponding gating strategies optimized for specific immunophenotyping tasks. Finally, the harmonization strategy implemented in ImmunoPheno enables reference databases to expand incrementally without reprocessing existing data, supporting scalable integration of public resources.

Together, these features enable ImmunoPheno to bridge large-scale single-cell reference data and conventional immunophenotyping experiments in a manner that is both scalable and directly actionable, extending beyond label transfer to inform how immunophenotyping experiments are designed, executed, and interpreted.

Our analyses also revealed general principles for applying ImmunoPheno in practice. While using reference data that closely matches the cytometry dataset generally improves annotation accuracy, we found that for small antibody panels (< 15 antibodies), it is often advantageous to also incorporate reference data from related tissues and alternative monoclonal antibodies targeting the same protein. Increasing the number of nearest neighbors used for annotation transfer helped mitigate noise arising from such differences. Finally, tuning parameters to balance sensitivity and specificity according to user needs was also important for achieving optimal performance.

Although ImmunoPheno is designed to be broadly applicable across immunophenotyping technologies and tissues, its performance ultimately reflects the scope and composition of the underlying reference data. Certain immune populations, such as granulocytes or cells defined primarily by intracellular markers, remain underrepresented in current proteo-transcriptomic datasets and are therefore more challenging to annotate or isolate using surface-marker-based approaches alone. In these settings, expert knowledge and complementary experimental strategies remain essential.

ImmunoPheno is thus not intended to replace expert-driven immunophenotyping, but rather to augment it by providing quantitative, data-driven guidance grounded in the collective knowledge encoded in large-scale single-cell datasets. By systematically leveraging existing data to inform antibody selection, gating strategy design, and cell identity annotation, ImmunoPheno aims to reduce subjectivity and improve reproducibility while preserving the flexibility required for expert interpretation and biological discovery.

More broadly, the ImmunoPheno framework is tissue-agnostic and can be used to construct analogous reference resources for other organs, tissues, and species in which standardized immunophenotyping strategies are less established. With the continued expansion of single-cell proteo-transcriptomic data, now numbering over 2,000 samples deposited in the NCBI Gene Expression Omnibus (GEO) repository, we anticipate that HICAR and other resources built using ImmunoPheno will continue to grow, enabling increasingly precise, quantitative, and reproducible immunophenotyping across a wide range of experimental contexts.

## Methods

### Normalization of protein expression data

For each antibody, we adapted a normalization strategy similar to that of Govek et al.^18^, fitting a statistical mixture model to the unnormalized expression levels pooled across cells. We consider two types of mixture models (Gaussian and negative binomial mixture models) with one, two, or three components, where the component with the lowest mean is assumed to correspond to the background.

The models are fitted using maximum likelihood estimation. For Gaussian mixture models, we use the expectation-maximization (EM) algorithm. For negative binomial mixture models, we use sequential least squares programming (SLSQP) to maximize the log-likelihood function. To facilitate convergence, the maximum likelihood estimates for the mean, variance, and mixture parameters of a Gaussian mixture model with the same number of components (fitted with the EM algorithm) are used as initialization for the binomial models. Once the one-, two-, and three-component models have been fitted, we use the Akaike Information Criterion (AIC) to identify the best-fitting mixture model.

Cells are then assigned to the background or signal components for each antibody. For one-component models, all cells are assigned to the background. For multi-component models, cells are classified as background if their expression value for the antibody is less than the mode of the background component.

Once all cells for an antibody 𝑝 have been classified, the expressions are normalized as,

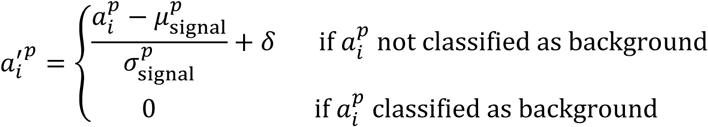

where 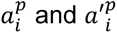 are respectively the raw and normalized expression values in cell 𝑖, 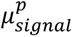 and 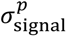 the mean and standard deviation of the raw expression values in cells not classified as background, and 𝛿 ≥ 0 a user-supplied parameter that sets the separation between the background 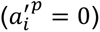 and the average of 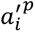 for cells expressing the protein.

To evaluate the normalization approach we considered published CITE-seq^37^, CyTOF^12^, and CODEX^38^ datasets of human PBMCs, BMMCs, and bone marrow tissue sections. These datasets contained panels of 114, 32, and 53 antibodies, respectively. We obtained the gene expression count matrices for three samples from the CITE-seq dataset (corresponding to titration levels 0.2x, 1x, and 2x), two samples from the CyTOF dataset (“benchmark dataset 2” in the original publication), and three samples from the CODEX dataset (samples “SB67_NBM27_H10_CODEX_Mesmer”, “SB67_NBM32_H37_CODEX_Mesmer”, and “SB67_NBM37_H35_CODEX_Mesmer”). We kept the original cell annotations from the CODEX and CyTOF datasets for downstream analyses, excluding cells labelled as “Artifact”, “Undetermined”, "Autofluorescent", or "CD44+ Undetermined" in the CODEX dataset, and “Not Assigned” in the CyTOF dataset. In the CITE-seq dataset, we filtered and annotated each sample independently based on the RNA gene expression data. Specifically, we used Azimuth^39^ with the following quality control filtering thresholds: min_nCount_RNA = 3,000, max_nCount_RNA = 10,000, min_nFeature_RNA = 1,000, and max_nFeature_RNA = 3,000, and excluded cell populations containing fewer than five cells. We converted all cell type identities to cell ontologies using the OBO Cell Ontology database^28^. We then normalized the protein expression data of each sample independently using the above procedure, where we used a negative binomial mixture model for the CITE-seq data, and Gaussian mixture models for the CODEX and CyTOF datasets. Cells expressing more than 80% of the antibodies in the panel (sig_expr_threshold = 0.80) were removed to reduce artifacts from non-specific binding, while those expressing fewer than 10% of the antibodies (bg_expr_threshold = 0.10) were excluded to filter out low-quality cells. We chose 𝛿𝛿 = 1.5 for the background-signal separation parameter. Antibodies that exhibited only background expression were removed. 2D visualizations were built using the UMAP Python package with default parameters.

### Implementation of the ImmunoPheno database

We implemented the database using MySQL due to its scalability, reliability, and public availability. Given the structured nature of the data, a relational model was chosen to organize information in a tabular format, ensuring consistency and facilitating complex queries. The database schema (Figure S1) was designed to split data into three primary categories: experiments, antibodies, and reference proteomic data. To ensure standardization throughout the database, antibody, antigen, cell type, and tissue names were mapped to Antibody Registry^26^, UniProtKB^27^, Cell Ontology^28^, and BRENDA tissue ontology^29^, respectively.

Data ingestion and database setup were automated using ImmunoPhenoDB, an auxiliary package that we developed to streamline the extraction, transformation, and loading (ETL) of harmonized datasets. This package provides MySQL procedures and functions to preprocess datasets and populate the database efficiently. Once populated, the database is hosted on an application server that exposes a public-facing REST API, enabling programmatic access to the stored data. The system design follows a microservices architecture (Supplementary Fig. 1B) to enhance scalability, flexibility, and performance. The core computational service was built using Flask, which was chosen for its lightweight design and seamless integration with Python libraries required for query execution. Given the computational demands of large-scale queries, Apache Spark was incorporated into the Flask service to parallelize MySQL queries, significantly reducing execution times.

To manage concurrent requests, Flask was deployed behind Gunicorn, which spawns worker processes to handle multiple connections. However, direct client access to Flask was restricted for security and performance reasons. Instead, a separate Express.js service serves as the primary REST API layer, acting as a request aggregator. To further optimize performance, Redis was integrated with Express to cache frequently requested data, reducing redundant database queries.

A NGINX reverse proxy sits in front of the Express service, providing load balancing and additional security measures such as request filtering and rate limiting. The entire system is containerized using Docker to ensure consistent deployment across environments and is hosted on an AWS EC2 instance for cloud-based accessibility and scalability.

To facilitate querying the database, we implemented a linear mixed-effects model to quantitatively assess the association between specific antibodies and cell populations, while accounting for potential batch effects across experiments. For each antibody, we used the MixedLM() function from the statsmodels Python library, modeling normalized protein expression levels as the dependent variable, cell type labels as the fixed effect predictor, and experiment ID as a random intercept effect. This allows the model to account for systematic shifts in signal intensity due to experiment-specific factors.

### Processing and harmonization of published proteotranscriptomic datasets for HICAR

We compiled a diverse set of publicly available single-cell CITE-seq and Ab-seq datasets across multiple tissues, including PBMCs, BMMCs, glioblastoma (GBM), and secondary lymphoid organs such as the spleen and lymph nodes. These datasets were selected based on their biological relevance, diversity of antibody panels, and tissue representation. To ensure standardization and reproducibility, we annotated each antibody in the datasets with their corresponding Research Resource Identifier (RRID). Antibodies for which we were unable to identify their RRID based on the information in the published paper were excluded from the dataset prior to ingestion into the database. Each dataset underwent quality control (QC) and protein normalization using ImmunoPheno to ensure consistency and harmonization across datasets.

Specifically, for the PBMC titrated dataset^37^, three titration conditions (0.2x, 1x, and 2x) were included. From the original 128-antibody panel, 14 antibodies were excluded: four isotype controls and ten antibodies due to being unable to find their RRIDs. Gene expression data was processed using Azimuth^39^. QC thresholds were set at nCount_RNA [3,000,1,000], nFeature_RNA [1,000, 3,000], with mitochondrial genes already filtered out per the original study. Protein expression normalization was performed across all conditions using ImmunoPheno with sig_expr_threshold = 0.80, bg_expr_threshold = 0.10, and bg_cell_z_score = 10. After normalization and filtering, 2,443 cells and 105 antibodies were retained.

For the PBMC dataset from individuals receiving influenza vaccination^58^, we considered 51 antibodies from the original 82-antbody panel, with 31 antibodies excluded due to being unable to find their RRIDs. Azimuth’s RNA QC thresholds were set at nCount_RNA [1,000, 3,500], nFeature_RNA [451, 999], percent_mt [0, 8]. Protein normalization used sig_expr_threshold = 0.80, bg_expr_threshold = 0.08, and bg_cell_z_score = 10. After normalization and filtering, 6,880 cells and 42 antibodies were retained.

For the PBMC dataset from COVID-19 patients^61^, we considered 153 antibodies from the original 192-antibody panel, with 39 antibodies excluded due to being unable to find their RRIDs. Azimuth’s RNA QC thresholds were set at nCount_RNA [3,000, 10,000], nFeature_RNA [1,000, 4,000], and percent_mt [0, 5]. Protein normalization used sig_expr_threshold = 0.80, bg_expr_threshold = 0.10, and bg_cell_z_score = 10. After normalization, 21,197 cells with 146 antibodies were retained.

For the dataset on infiltrating myeloid cells in glioblastoma^62^, we considered 251 antibodies from the original 268-antibody panel, excluding 17 antibodies (6 isotype controls and 11 for which we were unable to find their RRID). We used the same filtering and cell type annotations from the original study. Protein normalization used sig_expr_threshold = 0.80, bg_expr_threshold = 0.06, and bg_cell_z_score = 10. After normalization and filtering, 11,577 cells and 155 antibodies remained.

For the dataset on human bone marrow hematopoietic stem and progenitors^63^, we considered flow-through populations enriched for CD34^med/low^ cells. A total of 127 antibodies from the original 132-antibody panel were used, with 5 antibodies excluded due to being unable to find their RRIDs. We used the same filtering and cell type annotations from the original study. Protein normalized followed sig_expr_threshold = 0.80, bg_expr_threshold = 0.05, and bg_cell_z_score = 10. After normalization and filtering, 19,475 cells and 92 antibodies were retained.

For the PBMC dataset from psoriatic arthritis patients^64^, we considered 266 antibodies from the original 282-antibody panel, with 16 antibodies excluded (9 isotype controls and 7 antibodies for which we were unable to find their RRID). Azimuth’s RNA QC was performed with nCount_RNA [3,000, 20,000], nFeature_RNA [800, 3,000], and percent_mt [0, 10]. Protein normalization parameters were set at sig_expr_threshold = 0.80, bg_expr_threshold = 0.05, and bg_cell_z_score = 10. After normalization and filtering, 5298 cells and 154 antibodies were retained.

For the healthy bone marrow dataset^59^, we considered 22 antibodies from the original 25-antibody panel, with 3 antibodies excluded due to being unable to find their RRID. Azimuth’s RNA QC was set at nCount_RNA [1,000, 3,500], nFeature_RNA [400, 1,100], and percent_mt [0, 10]. Protein normalization followed sig_expr_threshold = 0.80, bg_expr_threshold = 0.10, and bg_cell_z_score = 10. After normalization and filtering, 5512 cells and 21 antibodies remained.

We also included a multi-tissue immune dataset spanning PBMCs, BMMCs, lung-associated lymph nodes, and spleen^65^. Of the 277-antibody panel, 261 antibodies were kept after removing 16 (nine isotype controls, seven without RRID). Azimuth’s RNA QC parameters were defined per tissue, with nCount_RNA [(3,000-4,000), (10,000-12,000)], nFeature_RNA [(1,200-1,500), (3,000-4,000)], and percent_mt [0, 15]. Protein normalization was applied consistently across tissues using sig_expr_threshold = 0.80, bg_expr_threshold = 0.05, and bg_cell_z_score = 10. After normalization and filtering, 12,263 cells and 146 antibodies were retained.

Finally, we incorporated an Ab-seq dataset profiling healthy and leukemic BMMCs and PBMCs^19^. All 197 antibodies from the original panel were used during processing. We used the same filtering and cell type annotations from the original study. Protein normalization parameters were sig_expr_threshold = 0.80, bg_expr_threshold = 0.10, and bg_cell_z_score = 10. After normalization and filtering, 14,246 cells and 197 antibodies remained.

### Generation and processing of CITE-seq data from human tonsils

Human tonsil tissue specimens were collected in sterile containers from immunocompetent healthy subjects undergoing routine tonsillectomy. The patients were a 23-month-old male, a 12-year-old male, and a 15-year-old female. Indications for tonsillectomy were obstructive sleep apnea or sleep disordered breathing. Written informed consent was obtained before surgery using a study protocol approved by the Children’s Hospital of Philadelphia (CHOP) IRB Committee. Approximately 1 gram of tonsil tissue was enzymatically digested into a single-cell suspension. The tissue was cut into 1 mm^3^ cubes and resuspended in 3 ml of prewarmed digestion media consisting of RPMI, 10% FBS, 1% PenStrep, 1% L-Glutamine, 1% Sodium Pyruvate, HEPES, 1000 units of recombinant DNase (Roche Diagnostics), 0.1 mg/ml liberase (Roche Diagnostics), and 1.6 mg/ml collagenase type 4 (Worthington Biochemical Corporation).

The tissue was then agitated on a 15° at angle using an orbital shaker rotating at 150 cycles per minute at 37 degrees. Every 10 minutes, the digestion media containing a single cell suspension was removed and replaced for a total of 4 cycles. Each single cell suspension was filtered using a 100 µm nylon mesh cell strainer and centrifuged (600 RCF, 10 minutes, 4°C).

Red blood cells were removed with lysis buffer (BD Biosciences) before further processing. The single cell suspension was then divided into three fractions. One fraction of 300 million cells was depleted of CD45+ cells using magnetic beads (MojoSort Human CD45 Nanobeads; Biolegend) to enrich for non-hematopoietic cells. A second fraction of 50 million cells was similarly depleted of CD19+ cells (MojoSort Human CD19 Selection Kit; Biolegend). A third fraction was left unmanipulated and therefore contained mostly B cells. A mixture of the three fractions (20% unmanipulated, 20% CD45 depleted and 60% CD19 depleted was divided into 5 million cell aliquots and cryopreserved in freezing media (90% FBS and 10% DMSO). Frozen aliquots were stored in liquid nitrogen until ready for sorting.

Cryopreserved cell suspensions were thawed in a water bath, diluted in 15 ml of complete media (RPMI with 20% FBS) and centrifuged (600 RCF, 5 minutes, 4°C). Pelleted cells were resuspended in 50 µl of cell staining buffer (Biolegend) and incubated with 5µl of Human TruStain FcX Fc Blocking reagent (Biolegend) for 10 minutes at 4°C. After 10 minutes, cells were stained with the TotalSeq-B antibody cocktail (Biolegend). Cells were then incubated for 30 minutes at 4°C. After 30 minutes, 5 ml of cell staining buffer was added, and the cell suspension was centrifuged (600 RCF, 5 minutes, 4°C). The cells were washed in 5 ml cell staining buffer two more times to remove residual antibodies. The cell suspension was then resuspended in 500 µl of cold PBS 1X buffer with 1% BSA and filtered through a 35 µm nylon mesh cell strainer. 1ug/ml of DAPI (Invitrogen) was added prior to sorting to label inviable cells. DAPI-negative live cells were sorted on a FACSAria Fusion Sorter (BD Biosciences) at The Children’s Hospital of Philadelphia Flow Cytometry Core.

To perform CITE-seq, the sorted cells were loaded onto Chip G for GEM generation and barcoding in accordance with the 10x Genomics protocol ‘Next GEM Single Cell 3’ kit v3.1 (dual index) with Feature Barcode technology for Cell Surface Protein’. Further cDNA amplification was performed, followed by the construction of gene expression libraries and cell surface protein libraries. Concentrations of cDNA and both libraries were measured using Qubit, and Agilent’s 2100 BioAnalyzer (High Sensitivity DNA Chip) was used to measure the fragment size and assess the quality. Final quantification was carried out using KAPA qPCR (Roche) and the libraries were sequenced using Novaseq6000 (Illumina) v1.5 paired-end 100-cycle sequencing kit (Read 1: 28 cycles, i7 index: 10 cycles, i5 index: 10 cycles, Read 2: 90 cycles). Cell Ranger was used to generate the output summary.

We processed the CITE-seq data of each individual sample using the same approach described in Section “Processing and harmonization of published proteo-transcriptomic datasets for the HICAR”, with nCount_RNA = [3,000, 20,000], nFeature_RNA = [1,200, 8,000], and percent_mt = [0, 13]. After normalization and filtering, 17,088 cells were retained. Protein normalization parameters were sig_expr_threshold = 0.80, bg_expr_threshold = 0.08, and bg_cell_z_score = 10.

### Imputation of protein expression data

Given a table of cells (rows) by antibodies (columns) containing reference protein expression data with missing values, we applied the following procedure to produce a complete table with no missing data.

First, we used the function KNNImputer from the scikit-learn Python library to impute missing values within each cell type. The number of neighbors was set to 10, or the number of cells in the cell population if fewer than 10 were present. Since missing values are not missing at random, some of the missing values are not imputable, and this procedure generally does not completely eliminate all missing values from the table.

To address the remaining missing values, we applied a greedy filtering procedure that removes one or more antibodies or cells at each step until no missing entries remain. Each row and column was assigned a weight based on the number of missing values it contained, scaled by a parameter 𝜌 ∈ [0,1]. Specifically, row weights were calculated as 𝜌 × (number of missing values), and column weights as (1 − 𝜌) × (number of missing values). The parameter 𝜌 (default 𝜌 = 0.5) controls the relative importance of rows (cells) versus antibodies (columns) in the filtering process. Higher values of 𝜌 prioritize the removal of rows with more missing data, while lower values favor removing columns.

We then iteratively removed the rows or columns with the highest weight, updating the weights at each step. If multiple rows or columns shared the highest weight, all were removed. When both rows and columns were tied in weight and count, we selected randomly between them.

To evaluate the imputation and filtering approach, we used complete, harmonized protein expression matrices from bone marrow^59^ and PBMC^58^ CITE-seq datasets, as well as our own tonsil CITE-seq dataset (see subsection “Generation and processing of CITE-seq data from human tonsils”). We assessed the accuracy of the k-nearest neighbor (KNN) imputation strategy (Fig. 4B) by simulating Missing Completely at Random (MCAR) scenarios, where a percentage of individual protein expression values were randomly masked. The masked dataset was then processed by our imputation algorithm. For values that remained missing after KNN imputation, we applied a multi-tiered fallback strategy: first, by random sampling from non-missing values for that antibody within the same cell type; second, by random sampling from non-missing values for that antibody across the entire dataset; and finally, by assigning a value of zero if the antibody was absent from all cells. Imputation accuracy was quantified using Lin’s Concordance Correlation Coefficient (CCC), comparing imputed values with their known ground-truth counterparts. This simulation was repeated ten times across varying levels of missing data (20%, 40%, 60%, and 80%).

To assess the efficiency of the greedy filtering algorithm in eliminating MNAR data (Fig. 4C), we simulated MNAR scenarios by systematically masking entire blocks of data corresponding to randomly selected subsets of cell populations and antibodies. This approach mimicked datasets in which specific markers or cell types were not measured. Following the application of our imputation and filtering algorithm, we quantified the fraction of the original data removed by the filtering step to yield a final, complete dataset. This simulation was repeated 100 times for each dataset.

### Automated design of antibody panels

We developed a computational approach to identify an optimal antibody panel of user-specified size k capable of effectively distinguishing target cell populations from background populations using reference data from the ImmunoPheno database. The workflow begins by retrieving protein expression data from the database based on user-defined filters such as target and background cell populations, tissues, and experiments. If no background is specified, all other cell types meeting the filters (excluding target cell types) are treated as background.

To reduce computational time, the resulting reference table is downsampled to a maximum of 10,000 cells. Missing values in the table are handled through the imputation and filtering procedure described in the subsection “Imputation of protein expression data”.

Classification and Regression Tree (CART) model is trained on the imputed table to classify cells as individual target populations or background. We use the DecisionTreeClassifier() implementation from the scikit-learn Python library. To enforce the exact number of desired antibodies, k, we apply a bisection search to identify the cost-complexity pruning parameter (cpp_alpha) that yields a tree with exactly k features. The resulting decision tree is further simplified by merging adjacent leaf nodes that predict the same class, eliminating redundant splits.

The selected antibodies are ranked by their feature importance score in the final CART model. Gating plots, along with the expected purity and yield, are then generated for each leaf node.

### Experimental validation of ImmunoPheno-generated antibody panels and gating strategies

#### MAIT cell and pDC isolation with FACS

Cryopreserved peripheral blood mononuclear cells (PBMCs; STEMCELL Technologies) were thawed in R10 medium (RPMI-1640 (Corning) supplemented with L-glutamine and 10% FBS (HyClone)) on the day prior to sorting and allowed to recover overnight under standard culture conditions (37°C, 5% CO2). On the day of sorting, cells were washed twice with PBS (Gibco) and then stained with Ghost Dye Violet 510 (Cytek Biosciences) for 30 min at 4°C in the dark to identify dead cells. Cells were then incubated in Human TruStain FcX (BioLegend) for 15 min at 4°C to reduce non-specific antibody binding. Cells were stained for 20 min at 4°C in the dark with one of the following antibody panels (all antibodies were from BioLegend unless otherwise indicated; Supplementary Table 2). For MAIT cell identification: PE/Cyanine7 anti-human CD4 (clone RPA-T4), PE anti-human CD161, and APC anti-human CD26 (clone BA5b) For pDC identification: FITC anti-human CD123 (clone 6H6), APC anti-human CD88 (C5aR), and APC/Cyanine7 anti-human CD45 (clone 2D1). Following staining, cells were washed with PBS supplemented with 2% FBS and 2 mM EDTA (Invitrogen), resuspended in PBS containing 1% FBS and filtered through a 35-μm mesh prior to sorting. MAIT cells and pDCs were sorted as viable CD4-CD26+CD161+ and CD45+CD123+CD88- populations, respectively, using a BD FACSDiscovery S8 cell sorter.

#### MAIT cell co-culture with THP-1 and activation

THP-1 cells (ATCC) were thawed and maintained in R10 medium supplemented with 0.05 mM 2-mercaptoethanol (Sigma-Aldrich) until cells grew as a uniform single-cell suspension. Isolated MAIT cells were co-cultured with THP-1 cells at a MAIT-to-THP-1 ratio of 1:5, with a minimum MAIT cell density of 1 x 10^4^ cells/mL, in a total volume of 100 µL/well in 96-well plates pre-coated with IFN-γ capture antibodies (Mabtech). Co-cultures were maintained under standard culture conditions for 12h prior to stimulation. MAIT cell TCR agonist, 5-OP-RU (TargetMol), was then added at a final concentration of 10 nM to stimulate the co-cultures, with DMSO (Cell Signaling Technology) used as the negative control. Following stimulation, cells were incubated for an additional 6h, after which activated IFN-γ secreting cells were quantified using an ELISpot assay.

#### pDC activation

Isolated pDCs were plated in 24-well plates at density of ≥1 x 10^5^ cells/mL in R10 medium supplemented with 10 ng/mL recombinant human IL-3 (Thermo Fisher). Cells were then stimulated with a TLR9 agonist, CpG-A oligonucleotide (InvivoGen), at a final concentration of 1µM and incubated overnight in standard culture conditions, with PBS used as negative control. Following stimulation, cells were transferred to 96-well plates pre-coated with IFN-α capture antibodies (Mabtech) such that each well contained ≥1 x 10^4^ cells in 100 µL of culture medium and incubated for an additional 6h. The frequency of IFN-α secreting cells was subsequently quantified using an ELISpot assay.

#### IFN-γ and IFN-α ELISpot assay

Prior to cell plating, ELISpot plates pre-coated with IFN-α and IFN-γ capture antibodies were washed five times with PBS and pre-incubated with R10 medium for ≥ 30 min. Cells were plated, stimulated and incubated as described above. After removing the cells, plates were washed with PBS five times, and 7-B6-1 biotinylated detection antibodies diluted in PBS containing 0.5% BSA (MilliporeSigma) were added at the manufacturer-recommended concentration and incubated for 2h at room temperature. Plates were then washed five times and incubated with Streptavidin-HRP diluted in PBS with 0.5% BSA for 1h.

Following five washes with PBS, color was developed by adding 100 µL TMB substrate solution per well and incubating until distinct spots emerged (5-10 min). The reaction was stopped by extensive washing of the plate and membrane underside with deionized water. Plates were dried overnight, and spots were counted manually under a microscope.

### Automated annotation of cell identities on immunophenotyping data

We adapted the STvEA^18^ algorithm for use within the ImmunoPheno platform. Given a query cytometry dataset, the system retrieves corresponding protein expression data from the ImmunoPheno server based on the monoclonal antibodies present in the query. To maximize the overlap with reference data, users can choose a more relaxed matching mode that relies on antibody targets rather than exact clone IDs. The retrieved reference data is then imputed using the method described in the subsection *“Imputation of protein expression data”* and used as the reference protein expression table for STvEA.

As described in Govek et al.^18^, STvEA employs a weighted, mutually nearest-neighbor approach within a consolidated space constructed from the normalized protein expression tables of the query and reference datasets. As part of its output, STvEA computes a transfer matrix that quantifies the contribution of each reference cell to the annotation of each query cell, normalized as a probability distribution over the reference cells.

To improve the annotation quality, we implemented several quality control filters to identify and remove low-confidence cell type assignments:

- *Distance ratio filter.* This filter evaluates the position of each query cell relative to its nearest reference cells in the consolidated protein expression space. We define the metric 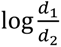, where 𝑑_1_ is the average distance from the query cell to its *k* nearest neighbors from the reference data, and 𝑑_2_ is the average pairwise distance among those neighbors. Query cell annotations with large distance ratios (outliers) are deemed unreliable and removed.

- *Entropy filter.* This filter assesses the uncertainty in each query cell’s annotation by computing the entropy of its label probability vector. Query cell annotations with large entropies (outliers) are deemed unreliable and removed.
- *Localization filter.* This filter assesses whether cells with the same cell type label cluster together in the protein expression space. We construct a cell-cell similarity graph based on the normalized protein expression data of the query dataset. In this graph, nodes are cells, and an edge connects two cells if their Pearson correlation exceeds the median correlation coefficient across the dataset. For each cell type, we generated a 2x2 symmetric contingency table classifying edges based on whether both, one, or neither cell belongs to the type. A Fisher’s exact test is used to evaluate enrichment of within-type edges. The FDR is controlled using the Benjamini-Hochberg procedure. Labels of cells assigned to non-significant (FDR > 0.05) cell types are removed.
- *Cell type merging filter.* This filter addresses the difficulty of distinguishing closely related cell types with similar protein expression profiles based on the antibody panel. We identify pairs of annotated cell types that are close (shortest path distance ≤ 2) in the OBO Cell Ontology. For each pair, we assess their separation in the protein expression space using a cell-cell similarity graph corresponding to cells from the two cell types and a Fisher’s exact test, as in the localization filter. Non-significant pairs of cell type annotations are merged into a single label. This process is repeated iteratively until no further merges are necessary.

### Evaluation of cell identity annotations

We established a comprehensive benchmarking framework to assess the accuracy of ImmunoPheno/STvEA single-cell identity annotations under a variety of conditions, including variations in antibody panel size and composition, the context of the reference data, and the criteria for matching antibodies between query and reference datasets. We selected two publicly available PBMC and bone marrow CITE-seq datasets^58,59^ to serve as query datasets for our evaluation. We utilized Azimuth^39^ to establish the identity of each cell based on its mRNA gene expression profile and used these labels as the ground truth for our evaluations. The protein expression data of both datasets were subsequently processed through the standard ImmunoPheno normalization and annotation pipelines described in subsections “*Normalization of protein expression data*” and “*Automated annotation of cell identities on immunophenotyping data*”. Antibodies with no detectable protein expression across all cells were excluded, resulting in a final panel of 22 antibodies for the bone marrow and 42 antibodies for the PBMC dataset. To investigate the impact of antibody panel size on annotation accuracy, we also simulated datasets with reduced antibody panels consisting of 10 antibodies. We employed a random forest classifier, as implemented in the function RandomForestClassifier from the scikit-learn Python library, with parameters n_estimators=100, max_depth=10, and min_samples_leaf=5, to rank all antibodies in each dataset based on their importance for predicting cell identity and select the top 10 antibodies in each dataset.

We then designed a series of benchmarks to test ImmunoPheno’s performance across three scenarios simulating different experimental contexts. As a baseline, we first used the query data itself as the reference, representing the theoretical best-case performance. The “same tissue” scenario restricted the reference data to all datasets in HICAR originating from the same tissue as the query, such as only PBMC datasets for the PBMC query, after excluding the query dataset itself. Finally, the “all tissues” scenario used the complete HICAR database, regardless of tissue origin, as the reference, again with the query dataset excluded. For each of these three scenarios, we assessed performance using both the full and reduced antibody panels and evaluated both strict clone ID and relaxed antibody target matching.

For all benchmarking tests, a consistent set of parameters was used for the annotation pipeline to ensure compatibility. Specifically, we set *ρ* = 0.6 for the imputation step, and k_find_nn = k_filter_anchor = 400 and k_transfer_matrix = 20 for the annotation step. Post-annotation filtering thresholds were determined dynamically based on the distance ratios and entropies generated within each benchmark.

The performance of all cell annotation tasks was quantified using MCC, providing a balance measure that is less susceptible to inflation by class imbalance. A challenge in comparing cell annotations is the varying level of granularity in cell type definitions (e.g., “B cell” vs. “memory B cell”). To ensure a fair comparison, we implemented a resolution-matching algorithm that leverages the hierarchical structure of the OBO Cell Ontology. When comparing a predicted label to a ground-truth label, if the predicted label is a direct descendant of the ground-truth label, it is considered a correct annotation at the resolution of the ground-truth. Conversely, predictions that are broader than the ground-truth, such as an ancestor, are not considered correct. This strategy requires ImmunoPheno annotations to be at least as specific as the ground-truth standard, penalizing overly broad or incorrect labels.

We benchmarked ImmunoPheno annotations against the NRN method and reference data from Triana *et al*.^19^. We used the PBMC and bone marrow reference datasets provided in the original publication for their method. We then compared the MCC scores obtained by ImmunoPheno, which leverages STvEA^18^ and the multi-experiment HICAR database, against those obtained by NRN, which uses a single proteo-transcriptomic dataset as its reference, using the same two query datasets as before, and evaluating both antibody clone ID and protein target matching strategies.

### Annotation of tumor-infiltrating immune cells from mIHC data from PDAC

Chromogenic mIHC images were generated using methods previously described^66^. 5 μm FFPE tissue sections from PDAC resections were stained with 22 antibodies delineating lymphoid and myeloid cell markers. Multiple 1 mm^2^ regions of interest (ROIs) were selected across the tumor specimen for single-cell characterization based on tissue quality post-staining and annotated in Aperio ImageScope (Leica Biosystems). Images were registered using the MATLAB Computer Vision Toolbox, AEC signal was extracted using FIJI^67^, and single-cell segmentation and labeling were performed with StarDist 2D^68^. The mean intensity for each marker was quantified in CellProfiler^69^, and gating thresholds for each marker were set in FCS Express Image Cytometry (DeNovo Software). Cell phenotypes were assigned based on a canonical immune phenotyping gating hierarchy and annotated in R statistical software.

We then annotated the same tumor immune cells using ImmunoPheno/STvEA. Protein 3-amino-9-ethylcarbazole (AEC) signal intensities were log-transformed (transform_type=”log”) and scaled (transform_scale=0.5) before fitting a Gaussian mixture model. Protein expression was normalized and filtered as described above, using parameters sig_expr_threshold = 0.90 and bg_expr_threshold = 0.10. Reference data were retrieved from HICAR through antibody target matching and imputed as described above, with imputation parameter 𝜌 = 0.25. ImmunoPheno/STvEA was then run with default parameters, except for k_transfer_matrix = 480 for PDAC1, and k_find_nn = 25, k_find_anchor = 10, k_filter_anchor = 25, and k_transfer_matrix = 320 for PDAC2. Post-annotation filtering thresholds were determined dynamically based on the distance ratios and entropies generated for each sample. After the annotation transfer and filtering, cell populations containing ≤20 cells were removed and replaced with “Not Assigned”. Expert and automated annotations were then compared using the MCC-based approach described in subsection *“Evaluation of cell identity annotations”*.

### Generation and analysis of CODEX data from human tonsils

Human tonsil tissue specimens were collected as described above. Tonsil tissue was submerged in RPMI media with 1% PenStrep and stored on ice until processed. Processing began by cutting two 2 mm blocks, which were placed in a tissue cassette, and submerged in a 4% paraformaldehyde solution (Electron Microscopy Sciences). The tonsil tissue was fixed for 24 hours at room temperature. After 24 hours, the tissue cassette was rinsed 3 times with PBS 1X for 10 minutes at 4°C before paraffin embedding by CHOP’s Pathology Core.

The CODEX panel consisted of 50 barcode-conjugated antibodies as well as the nuclear marker DAPI. Akoya antibodies were purchased preconjugated to their respective CODEX barcode. All other antibodies were custom-conjugated to their respective CODEX barcode according to Akoya’s PhenoCycler-Fusion user guide using the Antibody Conjugation Kit (Akoya, 7000009). CODEX staining was done using the Sample Kit for PhenoCycler-Fusion (Akoya, 7000017) according to Akoya’s PhenoCycler-Fusion user guide with modifications to include a photobleaching step in 4.5% H2O2 and 20 mM NaOH in PBS and overnight incubation in antibodies at 4°C. After staining, CODEX reporters were prepared according to Akoya’s PhenoCycler-Fusion user guide and added to a 96-well plate. The PhenoCycler-Fusion experimental template was set up for a CODEX run using Akoya’s PhenoCycler Experiment Designer software according to Akoya’s PhenoCycler-Fusion user guide. The PhenoCycler-Fusion experimental run was performed using Akoya’s Fusion 1.0.6 software according to Akoya’s PhenoImager Fusion User Guide. Images were captured on the PhenoImager Fusion microscope using default settings and subsequently pre-processed (stitching, registration, and background subtraction) by the Fusion software.

Image preprocessing and cell segmentation were performed by first generating composite nuclear and pan-membrane signals from the raw CODEX data. The nuclear signal was derived from the DAPI channel, while the pan-membrane signal was constructed by combining CD45, Vimentin, and Na+/K+-ATPase channels. Prior to combination, individual signal intensities were linearly rescaled between the 1st and 99.9th percentiles; the membrane signals were then summed and transformed using an exponential saturation function to ensure an asymptotic approach to the maximum range and minimize the influence of autofluorescence. Whole-cell segmentation was subsequently performed using the Mesmer model from DeepCell with an interior threshold of 0.3 and a maximum threshold of 0.2, and protein expression was quantified based on the total signal contained within the predicted segmentation masks. For quality control, the quantified data were analyzed as a Seurat object, where cells were filtered to remove outliers with counts above the 99.5th percentile, as well as cells with aberrant sizes below 25 pixels (∼6 um) or above 550 pixels (∼140 um).

We used ImmunoPheno to annotate the cell identities in the tissue section based on HICAR. CODEX protein expression levels were normalized using a Gaussian mixture model, as described in subsection “*Normalization of protein expression data*”, using parameters sig_expr_threshold = 0.80, bg_expr_threshold = 0.10, and bg_cell_z_score = 10. ImmunoPheno/STvEA was run with default parameters. The annotations were filtered as described in subsection “*Automated annotation of cell identities on immunophenotyping data*”, with parameters distance_ratio_threshold = 1.5 and entropy_threshold = 3.5.

To systematically map the spatial organization of the tissue, we performed a neighborhood analysis. Spatial proteomic data, including raw protein expression values, spatial coordinates, and cell annotations retrieved from ImmunoPheno, were imported into an AnnData object using the Scanpy package^70^. A spatial neighborhood graph was constructed using Squidpy^71^, connecting each cell to its 10 nearest spatial neighbors (sq.gr.spatial_neighbors, *k* = 10). Using this graph, we computed a neighborhood composition vector for every cell, representing the proportional frequency of each cell type within its immediate 10-cell neighborhood. We clustered neighborhood composition vectors using k-means clustering. To determine the optimal number of clusters, we computed the Davies-Bouldin Index using Scikit-learn for *k* values ranging from 5 to 30, identifying a global minimum at *k* = 16, which was selected as the optimal number of distinct spatial neighborhoods in this tissue.

## Code availability

The source code for ImmunoPheno is available at https://github.com/CamaraLab/ImmunoPheno (client) and https://github.com/CamaraLab/ImmunoPhenoDB (server). Documentation and tutorials are available at https://immunopheno.readthedocs.io/. An implementation of ImmunoPheno in Galaxy is available at https://toolshed.g2.bx.psu.edu/repository?repository_id=21c817fcdedbf487&changeset_revision=0cf5de252348. Docker images of ImmunoPheno are available at https://hub.docker.com/layers/camaralab/python3/immunopheno (client) and https://hub.docker.com/layers/camaralab/python3/immunophenodb (server).

## Data availability

Single-cell CITE-seq and CODEX count matrices, UMAP embeddings, and metadata from human tonsils have been deposited in Zenodo (https://doi.org/10.5281/zenodo.18405283). HICAR data are available through the ImmunoPheno website (https://immunopheno.org/) and via the ImmunoPheno Python package.

## Acknowledgements

We thank the Penn Cytomics and Cell Sorting Shared Resource Laboratory at the University of Pennsylvania (RRID:SCR_022376) for technical assistance with FACS during validation of ImmunoPheno gating strategies. We also thank Kathrin Brent and collaborators from the Childhood Hematopoiesis and Immune Development Atlas (CHIDA) project for making a subset of data available for illustrative purposes. This work has been supported by the U.S. National Institutes of Health (NIH) through grant U01CA269409 from the National Cancer Institute (NCI) Informatics Technologies for Cancer Research (ITCR) program (P. G. C.), as well as by the Pediatrics Networks for the Human Cell Atlas from the Chan Zuckerberg Initiative (P. G. C., K. T., N. R.), the Jeffrey Modell Foundation Translational Research Award (N. R.), and the grant R01AI179680 from the National Institute of Allergy and Infectious Diseases (NIAID) (N. R.).

## Author contributions

L. W. developed ImmunoPheno and performed all the computational analyses. M. A. N. carried out the experimental validation of antibody panels produced by ImmunoPheno. Z. Y. assisted with the implementation of ImmunoPheno. S. P. assisted with computational benchmarking analyses. S. S., N. K., and L. M. C. provided manually annotated mIHC PDAC datasets and assisted with their interpretation. A. J., K. J. A., J. S. T., E. C. C., N. R., and K. T. generated CITE-seq and CODEX data of human tonsils. P. G. C. and L. W. conceived the project and wrote the initial draft of the manuscript with contributions from the other authors.

