## Supplementary Figures and Tables for "ImmunoPheno: A Computational Framework for Data-Driven Design and Analysis of Immunophenotyping Experiments"

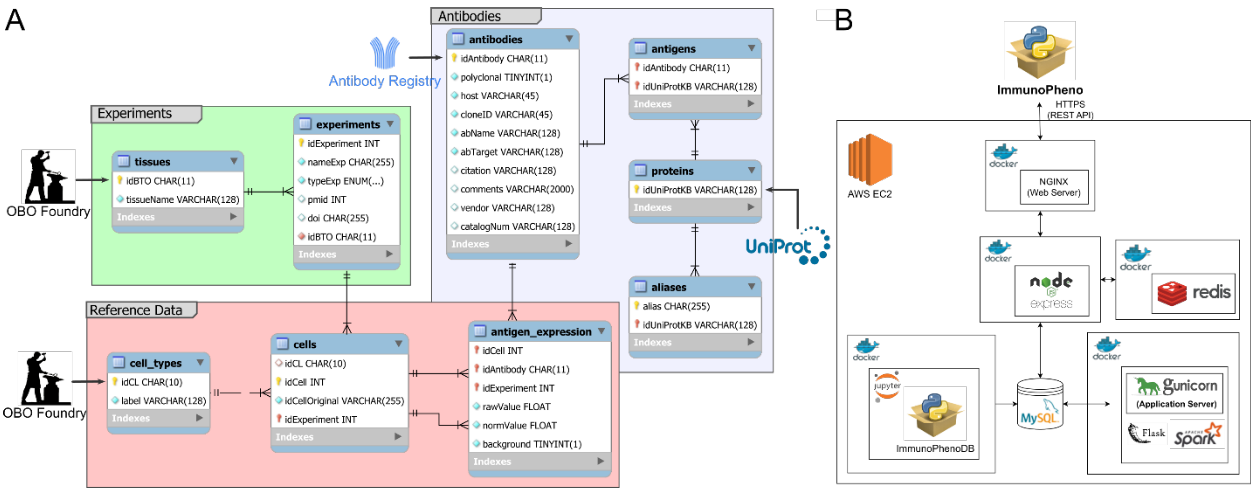


**Supplementary Figure 1. Architecture of the ImmunoPheno server. A)** Schema of the ImmunoPheno relational database. **B)** Modular and containerized structure of the ImmunoPheno server.

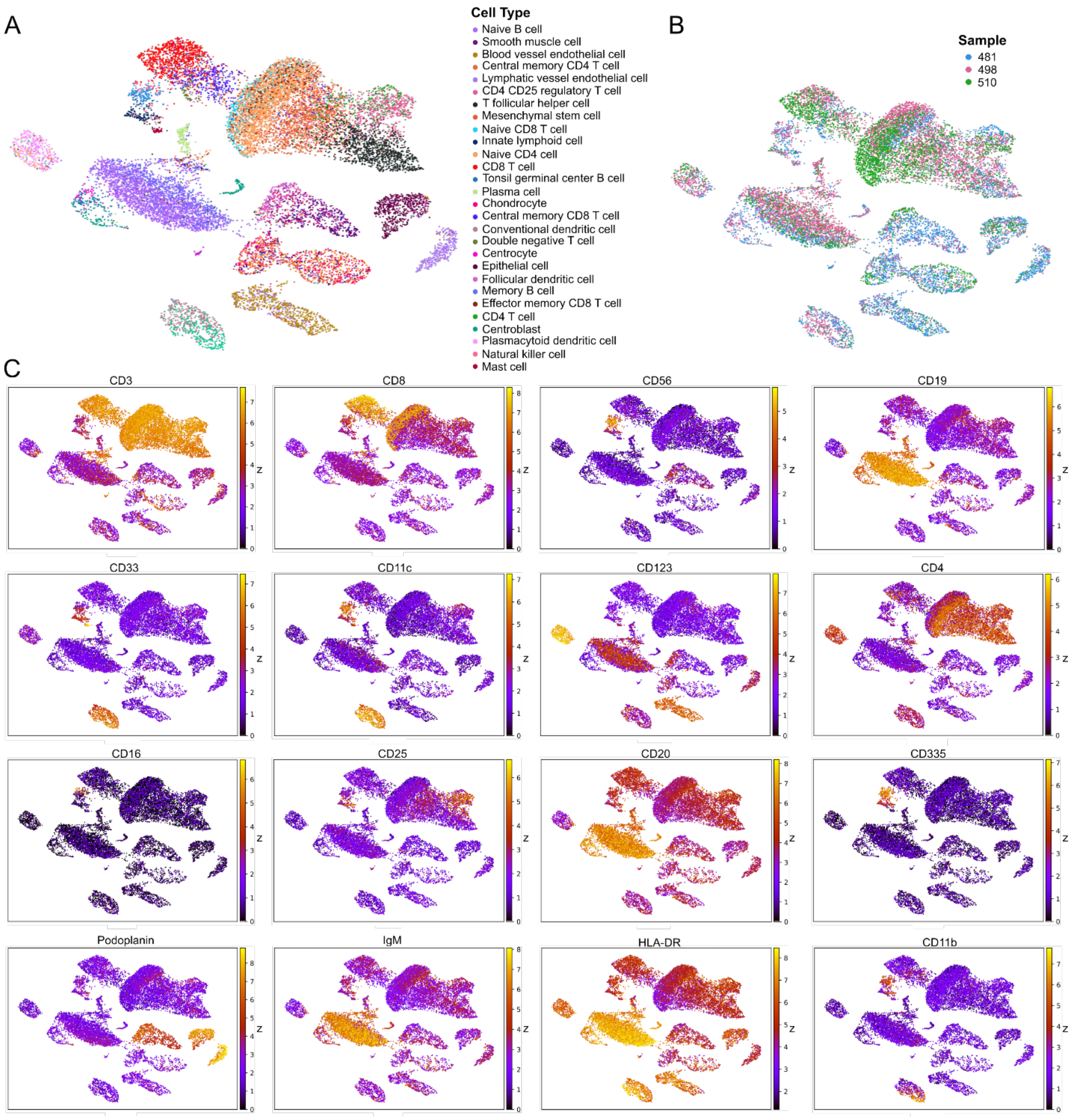


**Supplementary Figure 2. CITE-seq dataset of human tonsil using a 36-antibody panel. A)** UMAP representation of the mRNA data colored by the cell type annotations. **B)** UMAP representation of the mRNA data colored by the sample of origin. **C)** UMAP representation of the mRNA data colored by the normalized protein expression levels.

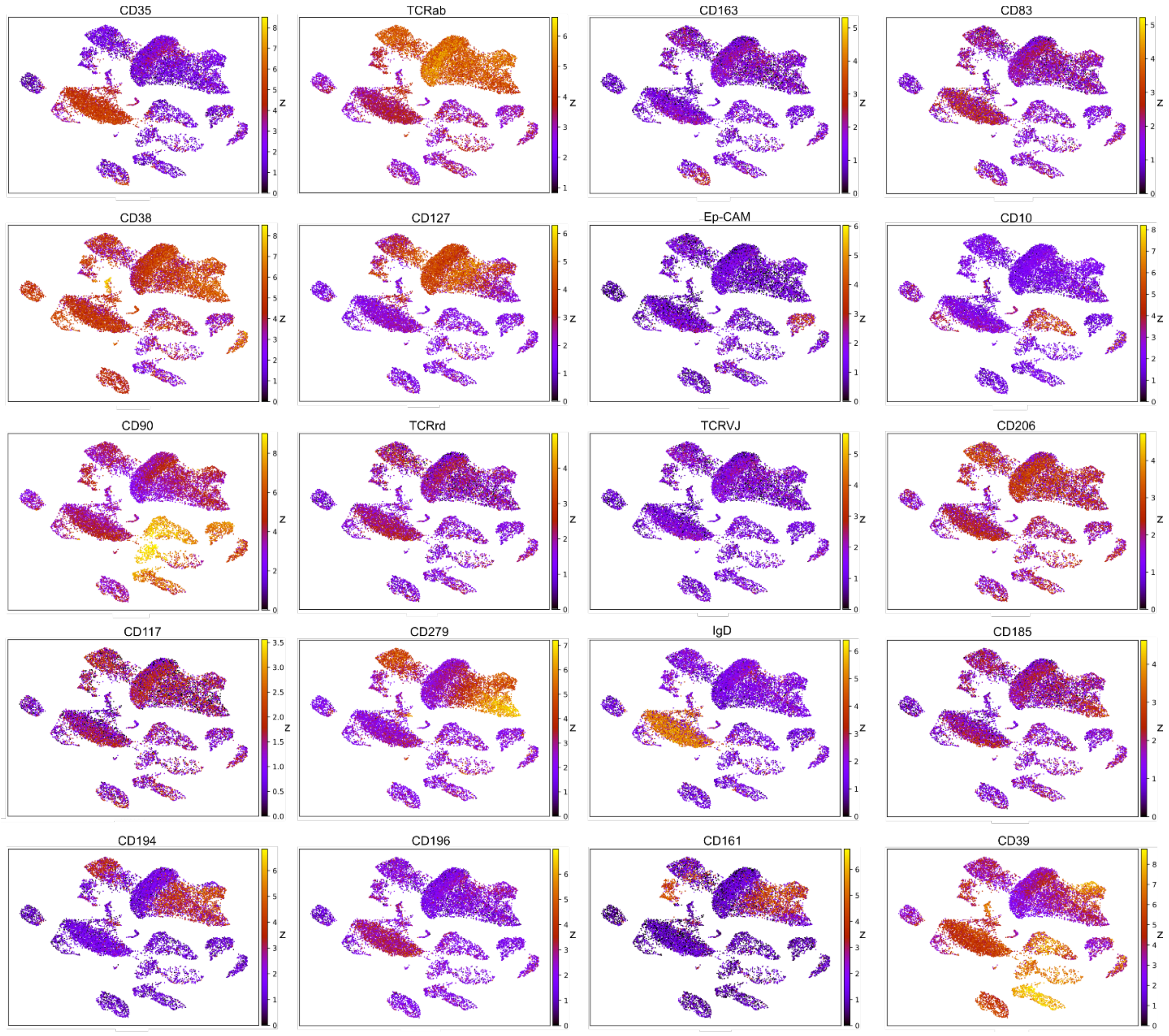


**Supplementary Figure 2C (continued).**

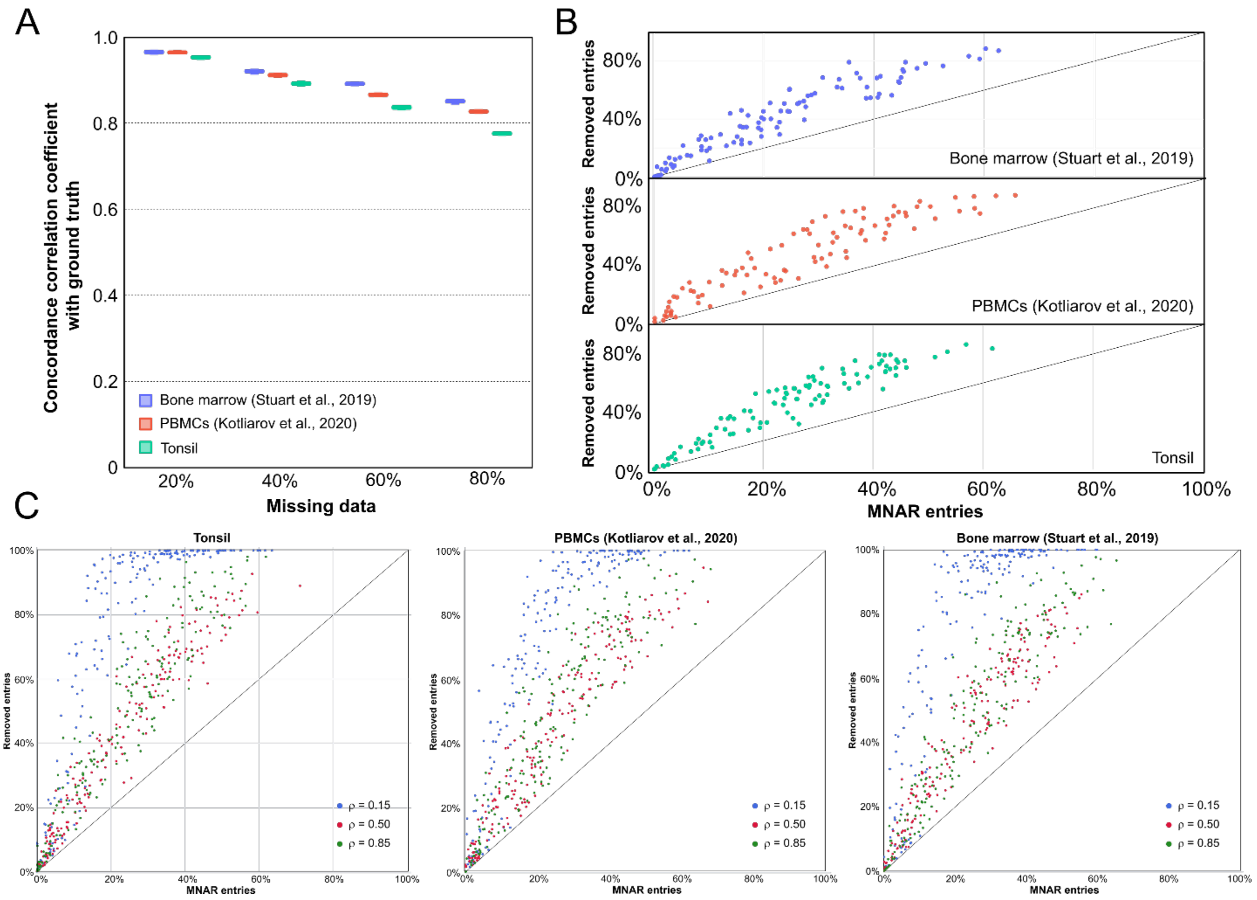


**Supplementary Figure 3. Cross-dataset protein expression imputation and filtering. A)** Concordance correlation coefficient between imputed and ground-truth protein expression values as a function of the percentage of missing data, shown across three datasets in which entries of the single-cell protein expression matrix were randomly masked. **B)** Percentage of data removed from the protein expression table by ImmunoPheno as a function of the percentage of simulated MNAR data, shown across the three datasets. **C)** Efficiency of post-imputation filtering as a function of the ρ parameter and the amount of MNAR data. Intermediate values of the ρ parameter (default ρ = 0.5), which balance the removal of cell populations and antibodies, generally yield the highest efficiency.

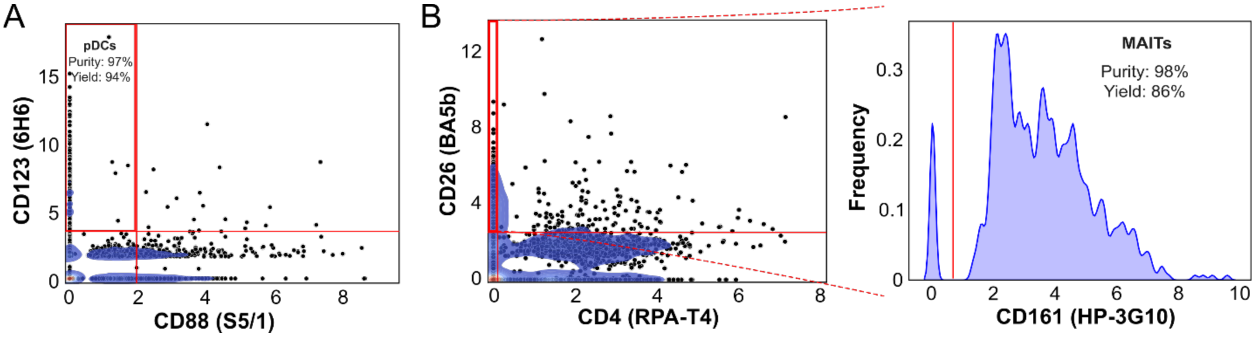


**Supplementary Figure 4. Small antibody panels and gating strategies for isolating rare immune cell populations generated by ImmunoPheno. A)** Two-antibody panel with gating strategy and expected yield and purity for isolating plasmacytoid dendritic cells (pDCs) from PBMCs. **E)** Three-antibody panel with gating strategy and expected yield and purity for isolating mucosal-associated invariant T (MAIT) cells from PBMCs.

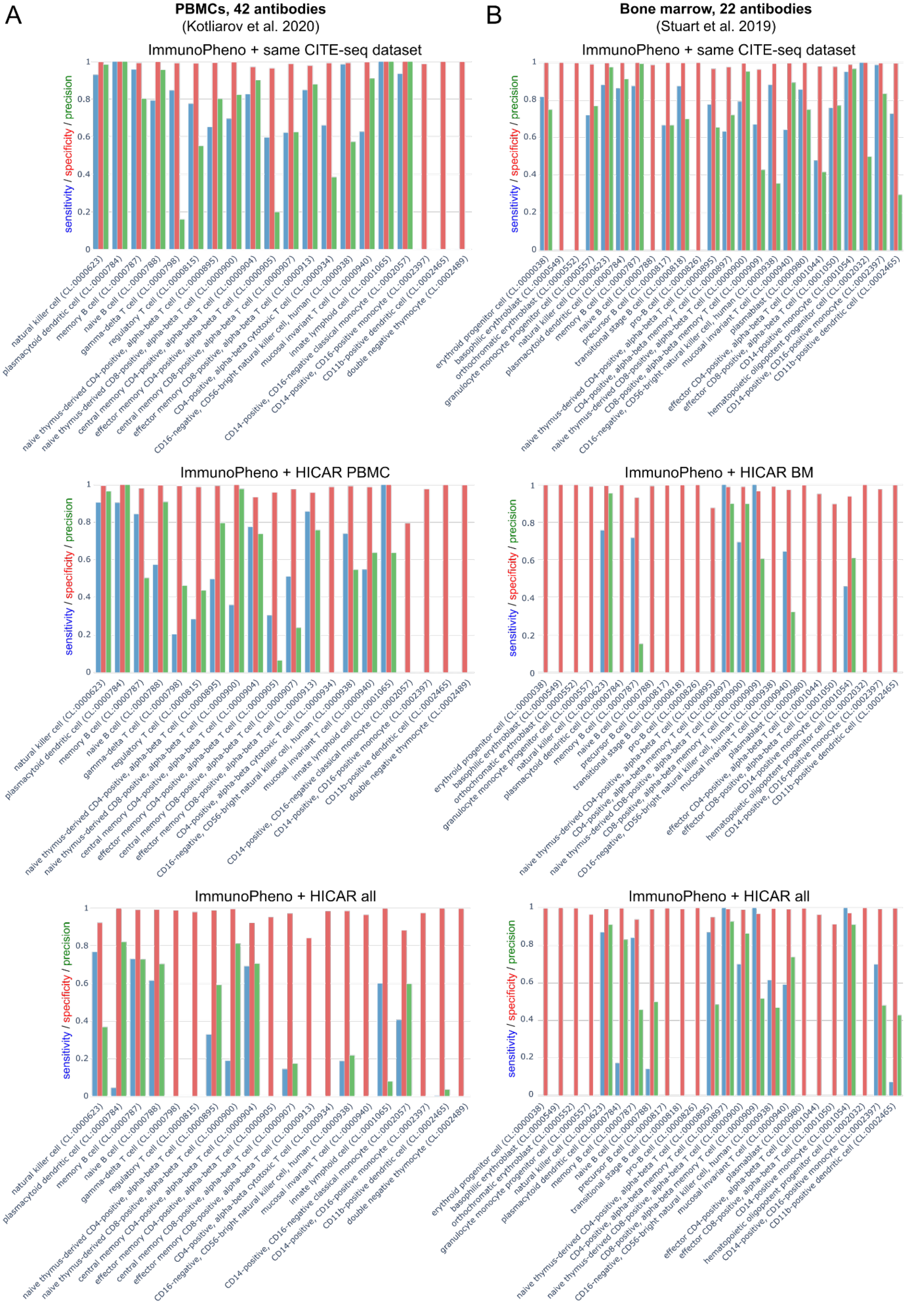
A
PBMCs, 42 antibodies
(Kotliarov et al. 2020)
ImmunoPheno + same CITE-seq dataset
sensitivity / specificity / precision
0
0.2
0.4
0.6
0.8
1
natural killer cell (CL:0000623)
plasmacytoid dendritic cell (CL:0000784)
memory B cell (CL:0000787)
naive B cell (CL:0000788)
gamma-delta T cell (CL:0000798)
regulatory T cell (CL:0000815)
naive thymus-derived CD4-positive, alpha-beta T cell (CL:0000895)
naive thymus-derived CD8-positive, alpha-beta T cell (CL:0000900)
central memory CD4-positive, alpha-beta T cell (CL:0000904)
effector memory CD4-positive, alpha-beta T cell (CL:0000905)
central memory CD8-positive, alpha-beta T cell (CL:0000907)
effector memory CD8-positive, alpha-beta T cell (CL:0000913)
CD4-positive, alpha-beta cytotoxic T cell (CL:0000934)
CD16-negative, CD56-bright natural killer cell, human (CL:0000938)
mucosal invariant T cell (CL:0000940)
innate lymphoid cell (CL:0001065)
CD14-positive, CD16-negative classical monocyte (CL:0002057)
CD14-positive, CD16-positive monocyte (CL:0002397)
CD11b-positive dendritic cell (CL:0002465)
double negative thymocyte (CL:0002489)
ImmunoPheno + HICAR PBMC
ImmunoPheno + HICAR all
B
Bone marrow, 22 antibodies
(Stuart et al. 2019)
ImmunoPheno + same CITE-seq dataset
erythroid progenitor cell (CL:0000038)
basophilic erythroblast (CL:0000549)
orthochromatic erythroblast (CL:0000552)
granulocyte monocyte progenitor cell (CL:0000557)
natural killer cell (CL:0000623)
plasmacytoid dendritic cell (CL:0000784)
memory B cell (CL:0000787)
naive B cell (CL:0000788)
precursor B cell (CL:0000817)
transitional stage B cell (CL:0000818)
pro-B cell (CL:0000826)
naive thymus-derived CD4-positive, alpha-beta T cell (CL:0000895)
CD4-positive, alpha-beta memory T cell (CL:0000897)
naive thymus-derived CD8-positive, alpha-beta T cell (CL:0000900)
CD8-positive, alpha-beta memory T cell (CL:0000909)
CD16-negative, CD56-bright natural killer cell, human (CL:0000938)
mucosal invariant T cell (CL:0000940)
plasmablast (CL:0000980)
effector CD4-positive, alpha-beta T cell (CL:0001044)
effector CD8-positive, alpha-beta T cell (CL:0001050)
CD14-positive monocyte (CL:0001054)
hematopoietic oligopotent progenitor cell (CL:0002032)
CD14-positive, CD16-positive monocyte (CL:0002397)
CD11b-positive dendritic cell (CL:0002465)
ImmunoPheno + HICAR BM
ImmunoPheno + HICAR all

**Supplementary Figure 5. Sensitivity, specificity, and precision of ImmunoPheno/STvEA single-cell identity annotations. A, B)** Sensitivity, specificity, and precision of the protein-expression-based annotations are shown for each cell type in two published single-cell CITE-seq datasets: PBMCs (A) and bone marrow (B). Ground-truth labels were derived from the mRNA gene expression profiles of the cells. Antibodies were matched between the reference and query datasets using antibody clone IDs. The tradeoff between sensitivity and specificity can be adjusted by modifying the ImmunoPheno/STvEA parameters.

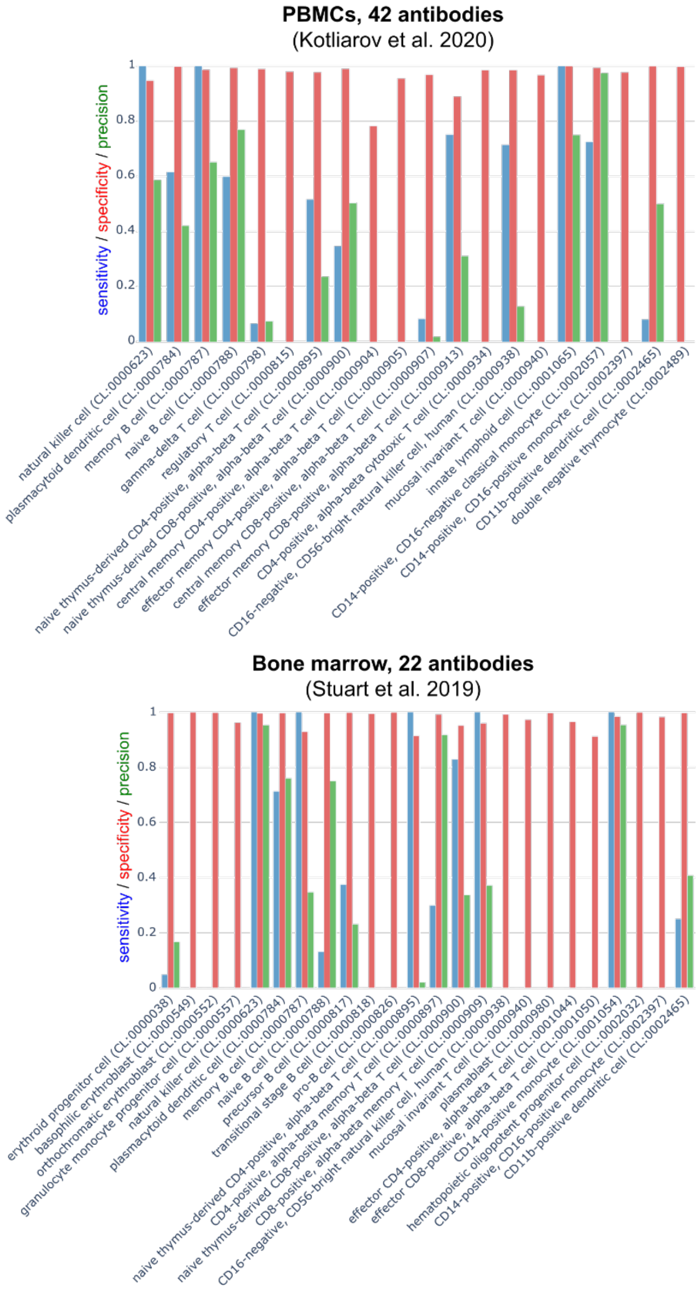


**Supplementary Figure 6. Sensitivity, specificity, and precision of NRN single-cell identity annotations.** Sensitivity, specificity, and precision of the NRN protein-expression-based annotations are shown for each cell type in the PBMC and bone marrow CITE-seq datasets of Supplementary Fig. 4. Ground-truth labels were derived from the mRNA gene expression profiles of the cells. Antibodies were matched between the reference and query datasets using antibody protein targets.

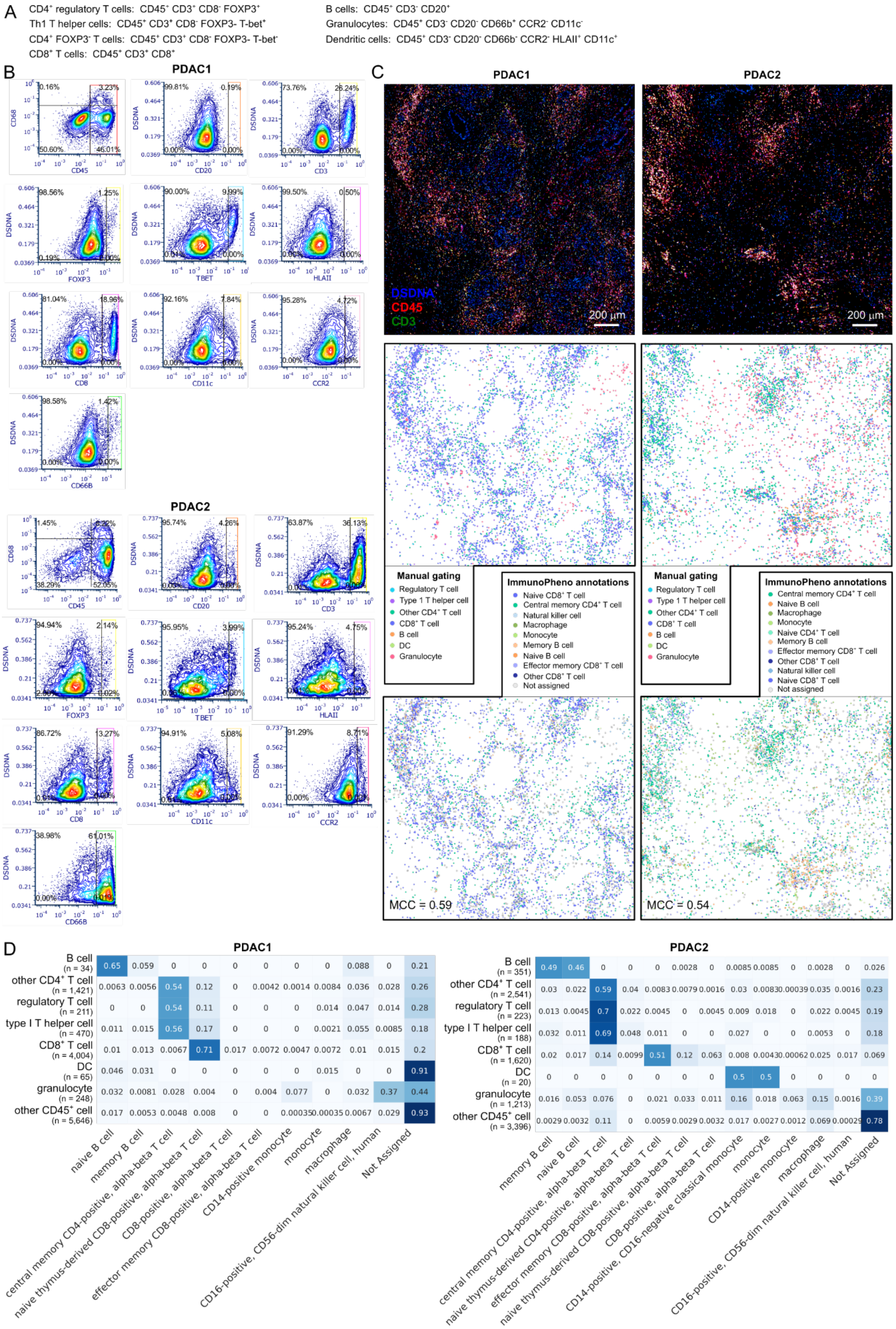
A
CD4+ regulatory T cells: CD45+ CD3+ CD8- FOXP3+
Th1 T helper cells: CD45+ CD3+ CD8- FOXP3- T-bet+
CD4+ FOXP3- T cells: CD45+ CD3+ CD8- FOXP3- T-bet-
CD8+ T cells: CD45+ CD3+ CD8+
B cells: CD45+ CD3- CD20+
Granulocytes: CD45+ CD3- CD20- CD66b+ CCR2- CD11c-
Dendritic cells: CD45+ CD3- CD20- CD66b- CCR2- HLAII+ CD11c+
B
PDAC1
PDAC2
C
PDAC1
PDAC2
DSDNA
CD45
CD3
200 µm
Manual gating
Regulatory T cell
Type 1 T helper cell
Other CD4+ T cell
CD8+ T cell
B cell
DC
Granulocyte
ImmunoPheno annotations
Naive CD8+ T cell
Central memory CD4+ T cell
Natural killer cell
Macrophage
Monocyte
Memory B cell
Naive B cell
Effector memory CD8+ T cell
Other CD8+ T cell
Not assigned
MCC = 0.59
MCC = 0.54
D
PDAC1
PDAC2

**Supplementary Figure 7. Annotation of immune cells in pancreatic ductal adenocarcinoma mIHC data. A)** Gating strategies used to define eight tumor-infiltrating immune cell populations. **B)** Application of these gating strategies to two tissue sections profiled with chromogenic mIHC. **C)** Top, tissue sections colored by three representative markers. Middle and bottom, cell annotations based on the gating strategies in (A) and ImmunoPheno's automated annotations of tumor-infiltrating $CD45^{+}$ cells, respectively. MCC: Matthews correlation coefficient between expert and ImmunoPheno single-cell annotations. **D)** Confusion matrices between manual and ImmunoPheno annotations.

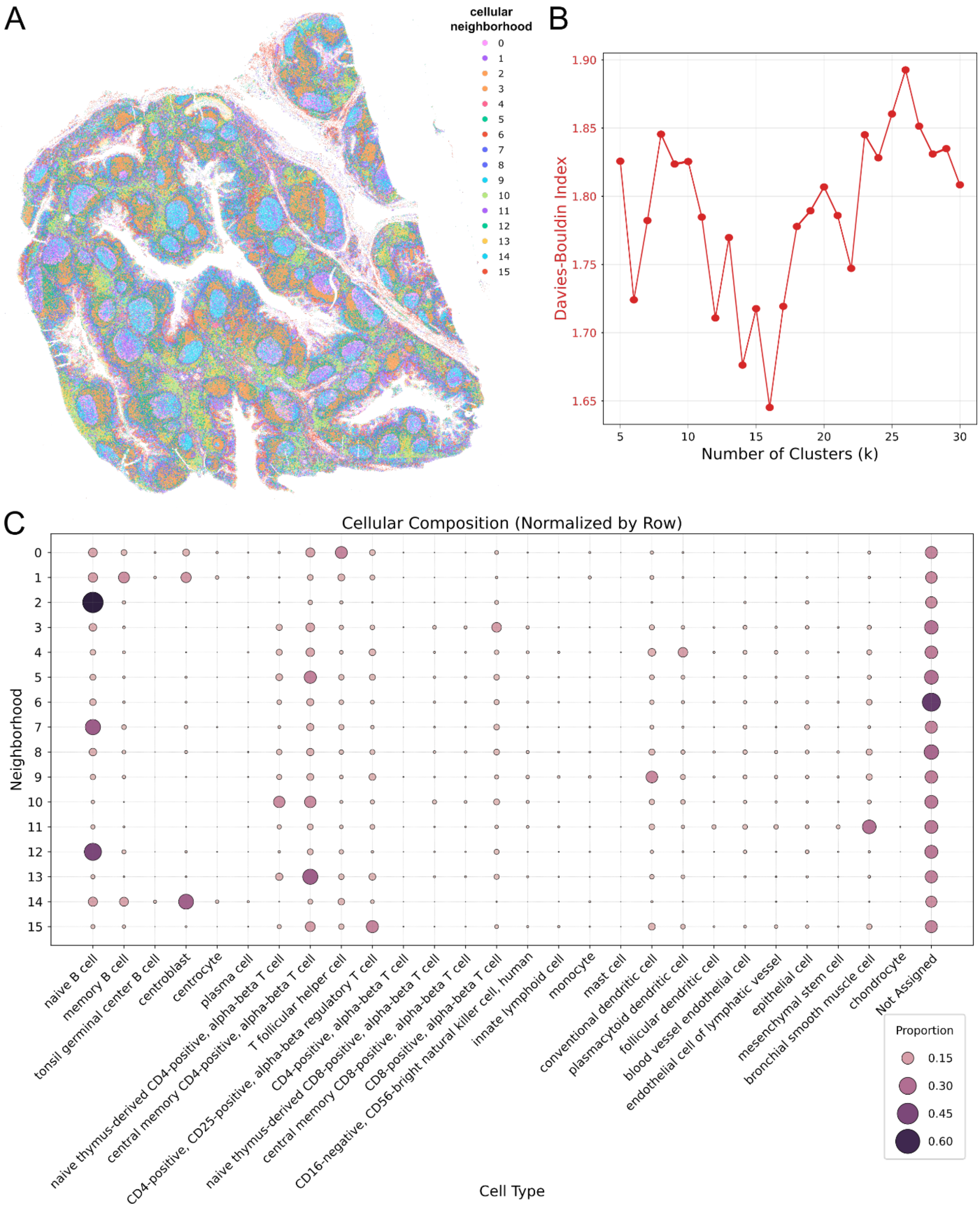


**Supplementary Figure 8. Neighborhood analysis of human tonsil tissue section annotated with ImmunoPheno. A)** Tissue section colored by the neighborhood identity of each cell. **B)** Davies-Bouldin clustering index for different values of *k* in *k*-means clustering, showing a minimum at 16 clusters. **C)** Cellular composition of each of the 16 cellular neighborhoods identified in the analysis.

### Supplementary Tables

| DOI | Description | Type | Cells | Antibodies | Tissue |
|---|---|---|---|---|---|
| 10.1038/s41598-022-24371-7 | PBMCs under titration | CITE-seq | 2,443 | 105 | Blood |
| 10.1038/s41591-020-0769-8 | PBMCs from influenza vaccination | CITE-seq | 6,880 | 42 | Blood |
| 10.1016/j.cell.2021.02.018 | PBMCs from COVID-19 patients | CITE-seq | 21,197 | 146 | Blood |
| 10.1038/s41593-020-00789-y | Myeloid cells infiltrating glioblastoma | CITE-seq | 11,577 | 155 | Glioblastoma |
| 10.1038/s41590-024-01782-4 | Bone marrow HSPCs | CITE-seq | 19,475 | 92 | Bone Marrow |
| 10.3389/fimmu.2022.835760 | PBMCs from psoriatic arthritis patients | CITE-seq | 5,298 | 154 | Blood |
| 10.1016/j.cell.2019.05.031 | BMMCs | CITE-seq | 5,512 | 21 | Bone Marrow |
| 10.1016/j.crmeth.2024.100938 | Immune cells across various tissues | CITE-seq | 12,263 | 146 | Blood, Bone Marrow, Lymph Nodes, Spleen |
| 10.1038/s41590-021-01059-0 | Healthy and leukemic BMMCs & PBMCs | Abseq | 14,246 | 193 | Blood, Bone Marrow |
| This paper | Immune cells from the tonsil | CITE-seq | 18,021 | 36 | Tonsil |

**Supplementary Table 1. Datasets in the Human Immune Cell Antibody Reference (HICAR).**

| Target | Fluorophore | Clone | Supplier | Cat. No. | Application |
|---|---|---|---|---|---|
| CD4 | PE/Cyanine7 | RPA-T4 | BioLegend | 300511 | MAIT cell isolation |
| CD161 | PE | HP-3G10 | BioLegend | 339903 | MAIT cell isolation |
| CD26 | APC | BA5b | BioLegend | 302709 | MAIT cell isolation |
| CD123 | FITC | 6H6 | BioLegend | 306013 | pDC isolation |
| CD88 (C5aR) | APC | S5/1 | BioLegend | 344309 | pDC isolation |
| CD45 | APC/Cyanine7 | 2D1 | BioLegend | 368515 | $CD45^+$ cell isolation |
| Viability | Ghost Dye Violet 510 | | Cytek Biosciences | 13-0870-T100 | Dead cell exclusion |
| Fc Receptor Block | | | BioLegend | 422302 | Non-specific binding reduction |

**Supplementary Table 2. Antibodies and dyes used for flow cytometry and cell sorting.**

| Tonsil CODEX antibody | Clone | Vendor | catalog number | RRID |
|---|---|---|---|---|
| CD31 | AKYP0047 | Akoya Biosciences | 4450017 | AB_2915935 |
| CD19 | MRQ-36 | Cell Marquee | custom | AB_3083084 |
| CD4 | AKYP0048 | Akoya Biosciences | 4550112 | AB_3094499 |
| BCL6 | IG191E/A8 | Biolegend | 648301 | AB_2274637 |
| CD196 | 53103 | Novus Biologicals | MAB195-100 | AB_2244231 |
| CD194 | L291H4 | Biolegend | 359402 | AB_2562364 |
| CD20 | AKYP0049 | Akoya Biosciences | 4450018 | AB_2915939 |
| CD117 | Polyclonal | Novus Biologicals | AF1356 | AB_354750 |
| Vim | RV202 | Novus Biologicals | NBP1-97672 | AB_2915945 |
| CD90 | D3V8A | Cell Signalling Technology | 16836 | AB_3076750 |
| CD68 | AKYP0050 | Akoya Biosciences | 4550113 | AB_293589 |
| CD279 | D4W2J | Cell Signalling Technology | 63815 | AB_367599 |
| RANKL | 12A668 | Novus Biologicals | NB100-56512 | AB_839050 |
| CD206 | Polyclonal | R&D Systems | AF2534 | AB_2063019 |
| FOXP3 | 236A/E7 | Invitrogen | 14-4777-82 | AB_467556 |
| CD45 | AKYP0074 | Akoya Biosciences | 4550121 | AB_2915946 |
| IGD | EPR6146 | Abcam | ab236778 | AB_3720828 |
| CD35 | E11 | Novus Biologicals | NBP2-33080 | AB_3283301 |
| CD11c | EP1347Y | Abcam | ab216655 | AB_2864379 |
| CD127 | A019D5 | Biolegend | 351302 | AB_10718513 |
| CD8 | AKYP0028 | Akoya Biosciences | 4250012 | AB_2915960 |
| CD161 | HP-3G10 | Biolegend | 339902 | AB_1501090 |
| VE-CAD | 16B1 | ThermoFisher Scientific | 14-1449-82 | AB_467495 |
| CD123 | IL3RA/2947R | Novus Biologicals | NBP2-79851 | AB_3080863 |
| CD183 | G025H7 | Biolegend | 353733 | AB_2563724 |
| CD25 | 4C9 | Cell Marquee | custom | AB_3720829 |
| CD26 | D6D8K | Cell Signalling Technology | 19775SF | AB_3720830 |
| HLA-DR | AKYP0063 | Akoya Biosciences | 4550118 | AB_3080864 |
| LYVE1 | EPR21857 | Abcam | ab232935 | AB_2889891 |
| ASMA | Polyclonal | Abcam | ab5694 | AB_2223021 |
| CD10 | MME/1892 | Abcam | ab237866 | AB_3080866 |
| CD185 | 51505 | Novus | MAB190-100 | AB_2292654 |
| CD11B | EPR1344 | Abcam | ab209970 | AB_2915959 |
| CD16 | D1N9L | Cell Signalling Technology | 72204SF | AB_3280014 |
| MADCAM1 | UMAB158 | ThermoFisher Scientific | UM500105CF | AB_3720831 |
| CD3e | AKYP0062 | Akoya Biosciences | 4550119 | AB_2936080 |
| TCR gamma/delta | H-41 | Santa Cruz Biotechnology | sc-100289 | AB_1130061 |
| IgM | MHM-88 | Biolegend | 314527 | AB_2563776 |
| Prox1 | EPR19273 | Abcam | ab236026 | AB_2894898 |
| CXCL12 | 79018 | R&D SYSTEMS | MAB350 | AB_2088149 |
| CD33 | 996810 | R&D SYSTEMS | MAB11371 | AB_2889385 |
| CD56 | MRQ-42 | Cell Marquee | custom | AB_3082973 |
| Na+K+ ATPase | EP1845Y | Abcam | ab167390 | AB_2890241 |
| CD163 | AKYP0114 | Akoya Biosciences | 4250079 | AB_2935895 |
| CD38 | AKYP0110 | Akoya Biosciences | 4250080 | AB_3082976 |
| CD79a | AKYP0109 | Akoya Biosciences | 4450078 | AB_3082977 |
| CD83 | HB15e | Biolegend | 305302 | AB_314510 |
| EPCAM | AKYP0119 | Akoya Biosciences | 4550088 | AB_2935888 |
| CD39 | AKYP0107 | Akoya Biosciences | 4250076 | AB_3096410 |
| PDPN | AKYP0007 | Akoya Biosciences | 4250094 | AB_3082979 |

**Supplementary Table 3. Antibody panel used for CODEX profiling of human tonsil tissue.**
